# A polymerase ribozyme increases copying fidelity through pyrophosphate-mediated RNA repair

**DOI:** 10.1101/2022.11.01.514781

**Authors:** Alexandra D. Kent, Lucy L. Yang, Kyle H. Cole, Ivee A. Lee, Andrej Lupták

## Abstract

Prior to the emergence of the contemporary biosphere, the first replicating systems are thought to have progressed through an RNA-based stage. Such an evolving world would likely have transferred heritable information during replication using RNA polymerase ribozymes. Though substantial effort has been put forth towards evolving RNA polymerases, many variants suffer from premature termination and low fidelity, resulting in low yields of full-length or active sequences. Replication of longer sequences requires a sufficiently high fidelity to lend an evolutionary advantage to an evolvable system. Here we demonstrate ribozyme-mediated repair of mismatched and damaged RNA sequences. Under conditions of saturating pyrophosphate concentrations, we show that a polymerase ribozyme can repair RNA sequences terminated in a mismatch, a non-extendable 2′-3′ cyclic phosphate, or both, to generate a triphosphorylated nucleotide. This repair step increases the fidelity and allows polymerization along an extended template, including the ribozyme itself. This increase of copying fidelity advances the longstanding goal of developing a self-replicating polymerase ribozyme.

## Main Text

Because the cellular functions of ribonucleic acids (RNAs) include critical roles in catalysis, translation, and metabolite-dependent regulation of gene expression, RNAs may have been the first biopolymers to store information and catalyze chemical reactions on an early Earth.^1–5^ The replication process is fundamental to life and is the mechanism by which heritable information is stored and propagated. In the extant biosphere, this is achieved through protein-based polymerase enzymes, but in a hypothetical RNA World—prior to the advent of genetically encoded proteins—replication would likely have been carried out by catalytic RNAs termed polymerase ribozymes.^6,7^ These ribozymes could serve as the central molecule linking the phenotype of an early protocell to its genotype, but sufficient activity and fidelity would be critical to build up an information storage system.^8,9^

The central reaction catalyzed by enzymes replicating phosphodiester-based informational polymers is the transfer of the *α*-phosphate of a triphosphorylated nucleoside to a 3′-hydroxyl of another nucleotide, with concomitant release of pyrophosphate (Fig. 1). The reverse reaction is pyrophosphorolysis of polynucleotides, yielding a nucleoside triphosphate and a one-residue shorter strand. Beginning with the first *in vitro* selection of a ligase ribozyme, a large body of work using molecular evolution and engineering has sought to isolate ribozymes that catalyze phosphodiester formation.^10–14^ The ligase ribozyme has been evolved into progressively more sophisticated polymerases capable of promoter-based polymerization, production of other functional nucleic acids, reverse transcription, and transcription of non-canonical (xeno-) nucleic acids, among other functions.^15–21^ However, most of the advanced polymerase ribozymes suffer from both mutation burdens that reduce the activity of produced transcripts and low yields of full-length products due to premature termination and degradation.^22–24^ Recent reports demonstrate that evolving the polymerase ribozymes further under selection pressure to produce active ribozyme sequences increased the fidelity enough to enable successive rounds of amplification of an active, short self-cleaving ribozyme.^25^ Even with these advances, replicating active polymerase ribozymes remains an unsolved challenge, highlighting a need for novel approaches to increasing replication fidelity. An early process for RNA repair could serve to increase replication fidelity, but very few mechanisms of RNA repair have been reported to date and none have been catalyzed by ribozymes. A mechanism for repairing both degraded and mismatched sequences would lend an evolutionary advantage to an evolving system and is an unexplored approach to increasing polymerase ribozyme fidelity.^26–28^

**Figure 1.**
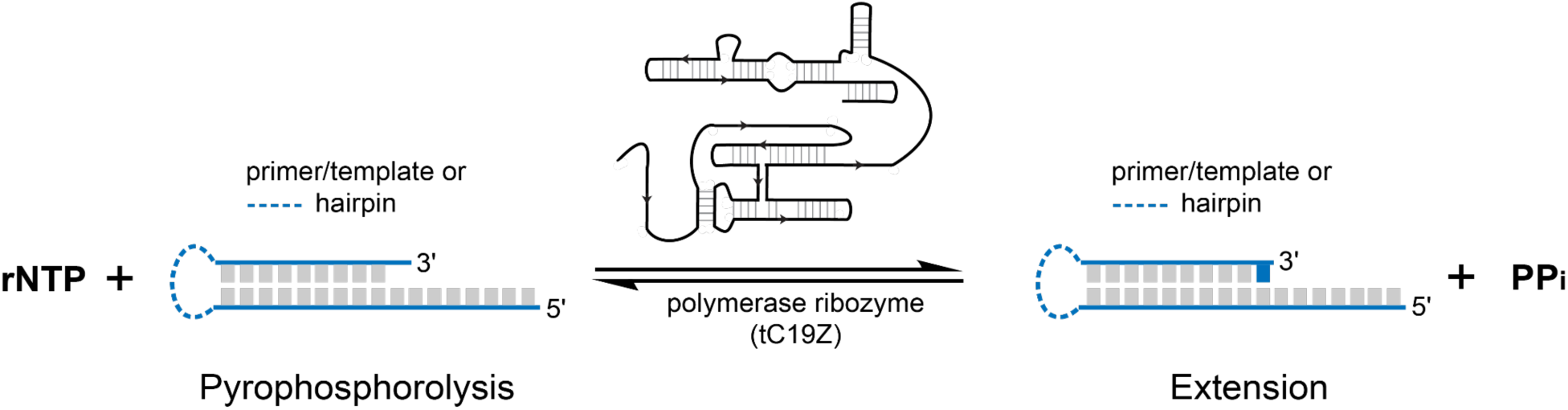
Reversible RNA synthesis by the polymerase ribozyme. The RNA polymerase ribozyme, tC19Z, catalyzes templated addition of a nucleotide to the 3′-end of a primer–template RNA with pyrophosphate (PPi) as a byproduct of the reaction. The reverse reaction, pyrophosphorolysis of RNA, removes the last nucleotide, yielding a nucleoside triphosphate. Dashed line indicates a linkage between the primer and the template, forming a single-molecule RNA hairpin used in this study.

We demonstrate that ribozyme-catalyzed pyrophosphorolysis of mismatched RNA sequences yields a nucleoside triphosphate, repairs the mutation, and increases the fidelity of the subsequent extension. Using sequencing analysis of mismatched and repaired RNAs, we show that repair of terminal mismatches improves downstream replication fidelity. Sequencing of RNAs generated from the polymerization of long templates reveal an increase in overall length and fidelity in the presence of pyrophosphate, and even include copies of the polymerase ribozyme. We further demonstrate that addition of pyrophosphate and the polymerase ribozyme to 2′-3′ cyclic phosphate-terminated (damaged) RNA sequences also facilitate damage repair and subsequent RNA polymerization. Thus, we present a mechanism for increasing the copying fidelity of long RNAs by a single polymerase ribozyme bringing us one step closer to a fully self-replicating RNA system.

## Results

One of the higher-fidelity RNA polymerases ribozymes, tC19Z, was selected *in vitro* from an earlier polymerase (R18), which in turn originated from the class I RNA ligase ^16,18^. The tC19Z sequence was evolved to have improved sequence generality and polymerase activity, although the template sequence also has a significant influence on replication efficiency. To extend a sequence, the 5′ end of the ribozyme base-pairs with the 5′ end of the template sequence and extends a primer (Fig. 1). Extension of the primer generates pyrophosphate, a byproduct of polymerization that has also been suggested as a source of available phosphate on an early Earth, which would have been critical due to the role phosphate plays in energy metabolism and genome replication.^29–33^ Inspired by a previous study that proposed pyrophosphorolysis was possible with a class I ligase derived ribozyme, we hypothesized that addition of pyrophosphate to the polymerization reaction would shift the equilibrium of the reaction and induce a low level of pyrophosphorolysis (Fig. 1) that could increase replication fidelity by selectively removing mismatched and damaged RNA.^34^

### Pyrophosphate-mediated mismatch repair

To maintain fidelity of genome copying at non-deleterious levels, modern replication machinery uses distinct active sites or entire protein complexes to correct mismatched base-pairs, but at the onset of evolvable systems, it is unlikely that mismatch repair was available as an independent biochemical activity.^35,36^ Thus, a simpler process for increasing replication fidelity would have been a desirable feature of an early replicating polymerase. To test whether ribozyme-mediated pyrophosphorolysis removes a mismatched nucleotide, we prepared hairpin sequences with 3′-terminal mismatches. We fused the primer and template to form a hairpin (Fig. 1) that serves both as a substrate for the tC19Z ribozyme at exact stoichiometry and as a practical construct for downstream analysis using high-throughput sequencing. We generated the hairpins *via* an *in vitro* transcription reaction of a self-cleaving HDV-like ribozyme (drz-Mbac-1) fused to the 3′ terminus of the target sequence.^37^ The ribozyme self-cleaves, yielding a 2′-3′-cyclic phosphate (2′-3′-cP) on the 3′ end of the primer segment. The cyclic phosphate can be removed enzymatically by T4 polynucleotide kinase (PNK) (Extended Data Fig. 1).^38,39^

We began by characterizing the reactivity of a terminal A/C mismatch in the primer/template sequence with the cyclic phosphate removed from the 3′ end. RNA polymerization by the ribozyme was not observed upon addition of nucleoside triphosphates (NTPs), indicating that an A/C mismatch acts as a chain-terminator (Fig. 2a,b). After addition of saturating concentrations of pyrophosphate (10 mM, with clearly visible Mg-PPi precipitate) and NTPs, we observed ribozyme-mediated extension of the sequence (Fig. 2), but we did not observe the −1 product by denaturing polyacrylamide gel electrophoresis (PAGE), likely because the equilibrium highly favors polymerization (SI Note 1, Fig. 2b, and Extended Data Fig. 2). When removed through pyrophosphorolysis, the terminal nucleotide is converted into a nucleoside triphosphate, and an A/C-terminated hairpin would produce a molecule of ATP. To obtain independent evidence of pyrophosphorolysis, we detected this critical metabolite using an ultrasensitive firefly-luciferase-based assay, which yields a photon for each ATP molecule present in the reaction.^40^ Because pyrophosphate inhibits the luciferase reaction, insoluble pyrophosphate was first removed from the samples by centrifugation, and the remaining pyrophosphate was subsequently hydrolyzed by addition of pyrophosphatase enzyme. The resulting samples were incubated with the firefly luciferase and D-luciferin and were then imaged using an ultracooled EM-CCD camera. The integrated luminescence was measured for both the pyrophosphorolysis reaction and control reactions lacking one of the components. In the absence of either pyrophosphate, the ribozyme, or the hairpin substrate, we observed a baseline level of luminescence. In reactions where all components were present, the luminescence was significantly greater, indicating pyrophosphate-mediated synthesis of ATP (Fig. 2c and Extended Data Fig. 3). These experiments provide evidence that the mechanism of repair occurs by a pyrophosphorolysis reaction and demonstrate the first instance of ribozyme-catalyzed production of ATP.

**Figure 2.**
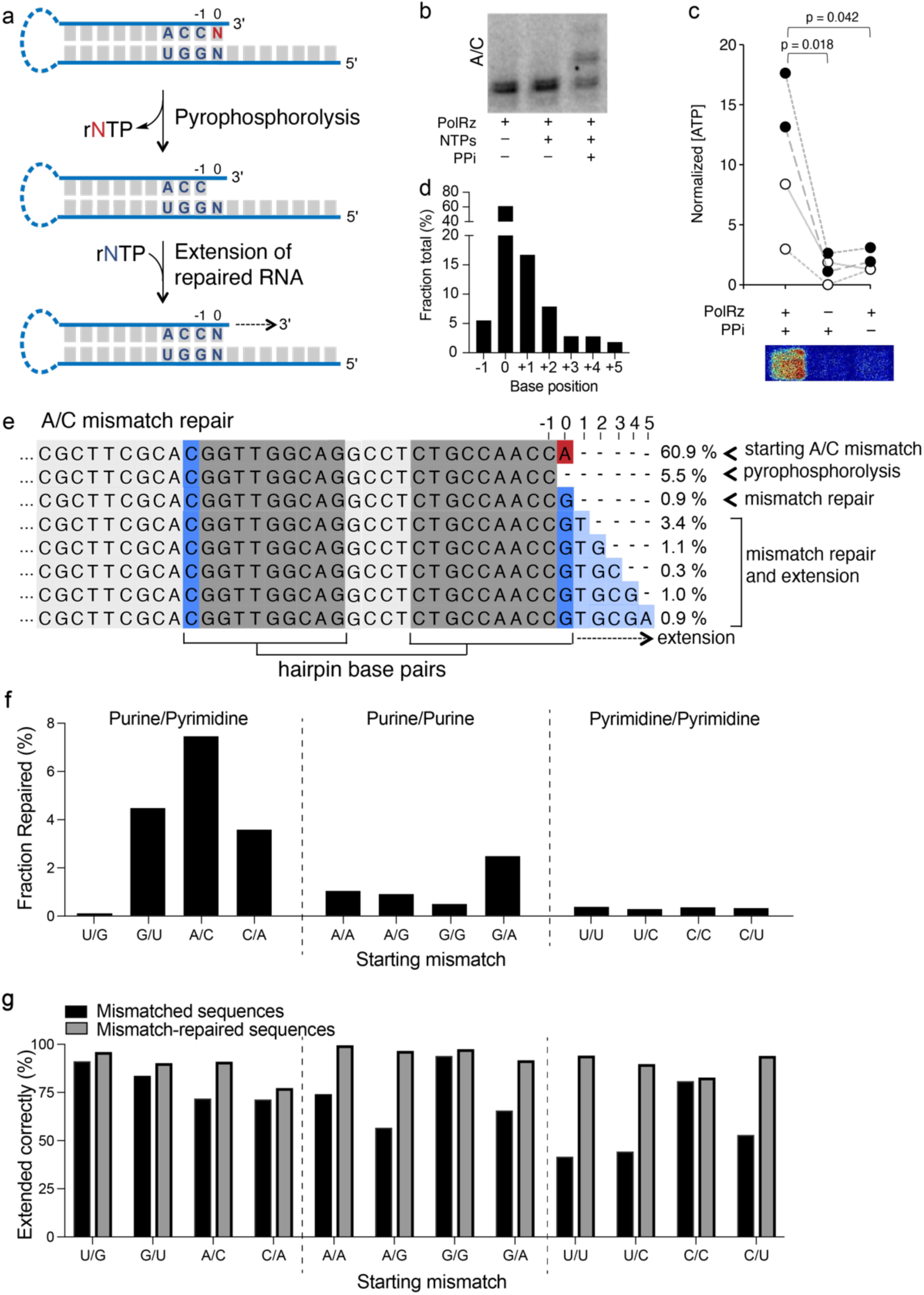
Pyrophosphorolysis-mediated mismatch repair and ATP generation by a polymerase ribozyme. (a) Graphical representation of a 3′ terminal mismatch (shown in red) undergoing ribozyme-catalyzed pyrophosphorolysis and subsequent repair. (b) High-resolution PAGE analysis of a ^32^P-labeled A/C mismatched hairpin. The left two lanes show no extension of the hairpin in the absence of nucleotides or pyrophosphate. Upon addition of pyrophosphate, the sequence can be extended (right lane). (c) ATP generated by the removal of the terminal AMP of an A/C-mismatched hairpin. Reactions lacking either the polymerase ribozyme or pyrophosphate show only background levels of luminescence by firefly luciferase. Open and filled circles represent two independent RNA preparations. Gray lines link experiments performed side-by-side from the same component stocks. Representative luminescence image is shown below. (d) Distribution of sequences extended by the ribozyme in the presence of pyrophosphate (mismatched and repaired), determined by HTS. (e) HTS analysis of sequences of the A/C-mismatched hairpin after incubation with the polymerase ribozyme and PPi. The sequences show the starting A/C mismatch (red A), the removal of the mismatched nucleotide by phosphorolysis (second row sequence, comprising 5.5% of total hairpin reads), the addition of the correct nucleotide (blue G at position 0), and the extension of the corrected sequence (light-blue boxes at positions 1–5). Total percentage of sequences extended are shown in d. (f) Fraction of each mismatch repaired through ribozyme-catalyzed pyrophosphorolysis, grouped by base-pair type. (g) Fidelity of extensions, comparing the fractions of the correctly extended sequences in sequences containing a mismatch (black bars) and the correctly extended sequences that are extended after mismatch repair (gray bars), indicating the fidelity of all sequences that follow the mismatch-repaired position. In all cases, pyrophosphate-mediated mismatch repair leads to an increase in the fidelity of extended sequences. The large differences between the dark and light bars imply that unrepaired mismatches lead to further errors in RNA polymerization.

To further analyze the repair reaction, we used high-throughput sequencing (HTS) of the mismatched-terminated hairpin reactions (Fig. 2d,e). HTS analysis revealed that pyrophosphorolysis of the substrate yields approximately 100-times higher concentrations of a hairpin lacking the terminal nucleotide than the no-pyrophosphate controls (Extended Data Fig. 4), providing experimental confirmation for the pyrophosphorolysis of the terminal nucleotide.

We repeated this analysis for each of the eleven other possible combinations of terminal mismatches (Fig. 2 and Extended Data Figs. 2, 4, and 5). Sequence analysis of the extension reactions correlated well with the gel electrophoresis analysis. In addition, the HTS data provided a deeper insight into the reactions, revealing the fractions of the sequences that were repaired and subsequently extended. PAGE analysis was consistent over replicate experiments. We found that the efficiency of repair and extension depends on the identity of the mismatch: purine/pyrimidine mismatches were repaired the most efficiently, with the exception of the U/G wobble. Purine/purine mismatches were repaired and extended at a low frequency upon the addition of pyrophosphate but were not extendable by the ribozyme in the absence of pyrophosphate (Fig. 2f,g and Extended Data Fig. 5). In contrast, pyrimidine/pyrimidine mismatches were extended well by the ribozyme, but most of the extension products retained the mismatch (Fig. 2f,g and Extended Data Fig. 5). Sequences containing terminal C/A or G/U mismatches behave like the A/C mismatch and only showed extension (with varying efficiencies) after pyrophosphorolysis of the mismatch.

In addition to PAGE analyses, we obtained quantitative results for the fidelity of extension products from the sequencing data. We measured the fidelity at each position of the extension and found that for all mismatches, repair of the mismatched base led to an increase in the fidelity of downstream extensions (Fig. 2f,g). In some cases, such as the pyrimidine/pyrimidine mismatches, the effect is strongly pronounced: although the frequency of repair is low, when a hairpin is repaired, the extension occurs with high fidelity. Specifically, in the case of the U/C mismatch, extension of the terminal mismatch preferentially adds a second mismatch (C/A) which then acts as a chain terminator (Extended Data Fig. 6). Therefore, we conclude that mismatched sequences that do not terminate the extension are more likely to acquire additional mutations, as has been observed in enzyme-free polymerization of nucleic acid sequences, implying a mechanism for error propagation that must involve an allosteric modulation of the fidelity of incoming nucleotide incorporation by upstream mismatches.^41^

We observed that addition of pyrophosphate inhibited the forward polymerization reaction in the positive control (Supplementary Fig. 1). To quantify the pyrophosphate reaction inhibition, we used a non-hairpin, primer−template complex to measure reaction kinetics of the forward extension reaction. Initial rates were plotted as a function of substrate concentration and revealed an approximate K_M_ of 4 mM with respect to NTPs (Extended Data Fig. 7A,B,C). We then measured the reaction kinetics in the presence of pyrophosphate and plotted the initial rates of tC19Z-mediated polymerization as a function of pyrophosphate concentration (Extended Data Fig. 7D,E). At millimolar PPi concentrations, the forward reaction was inhibited approximately five-fold, compared to when no PPi is present.

### Addition of pyrophosphate to extension reactions of a long template results in increased length and fidelity

Based on our demonstration that a polymerase ribozyme can repair hairpin sequences terminating in a mismatch and the mechanism for that repair, we hypothesized that pyrophosphate-mediated repair would suppress mutations in long extension reactions of primers, leading to increased fidelity in the polymerization of long templates. To do this, we used a primer–template system that can result in a templated polymerization of 118 nucleotides (Fig. 3a). We performed the ribozyme–catalyzed primer extension reactions for either three hours or four days at varying concentrations of pyrophosphate, and the RNAs were analyzed using HTS. As the concentration of pyrophosphate increased, we observed an increase in the average length of the primer extensions, although the total number of sequences decreased overall (Fig. 3b, Supplementary Table 3), consistent with pyrophosphate inhibition of polymerization. As hypothesized, the sequencing data showed an increase in fidelity at saturating concentrations of pyrophosphate (Fig. 3c,d). Further analysis revealed that many sequences resulting from early termination had mismatches at the 3′-termini, suggesting that the mismatches caused the polymerization to abort (Supplementary Fig. 2). We observed a negative correlation between the number of sequences and their length in the three-hour experiment, suggesting a trade-off between copying fidelity and the number of extension products. We also observed complements of tC19Z sequences in the HTS data with strand-specific ligation products from sequencing library preparation, indicating that the sequences originated from off-target priming and extension of the polymerase ribozyme itself. Nearly full-length (98%) ribozyme sequences were present in samples containing pyrophosphate and exhibited higher fidelity, whereas no long-read RNA segments antisense to the ribozyme were detected in the no-pyrophosphate HTS dataset (Supplementary Fig. 3).

**Figure 3.**
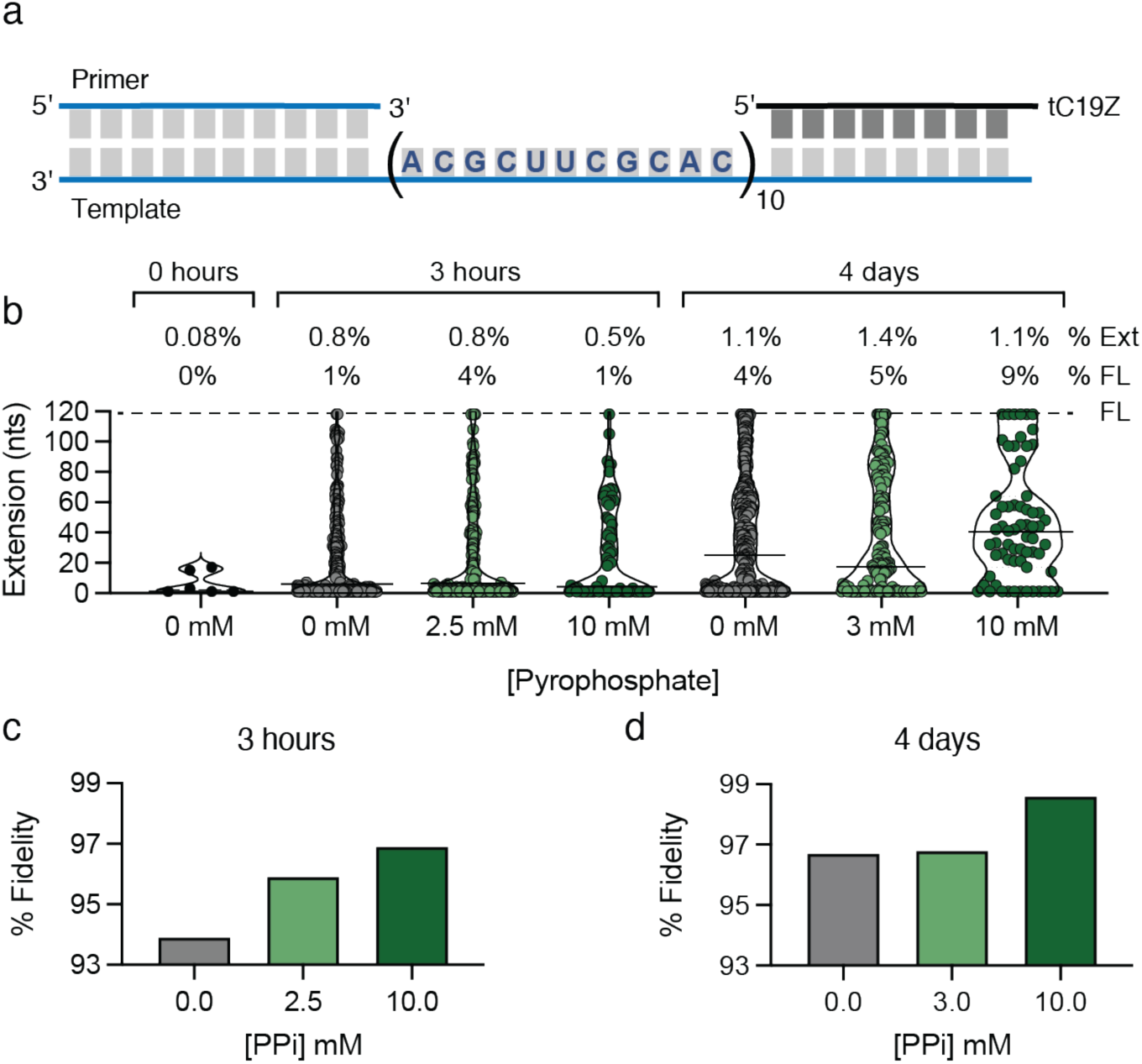
Effect of pyrophosphate on RNA polymerization along a long template. (a) The primer–template construct used for long extension experiments. (b) Plots showing the length of sequences resulting from polymerase-mediated extension of the primer at varying concentrations of pyrophosphate at either three hours or four days. Black horizontal bar indicates average length of sequences. FL dashed line indicates full length sequences. “% Ext” shows the percentage of sequences that were extended from primers, and % FL indicates the percentage of extended sequences that are full length. (c,d) Graphs showing the overall fidelity of extension products at varying concentrations of pyrophosphate at either three hours or four days. All analyses are based on HTS data. At ∼10 mM, Mg-PPi is above saturating concentration and is partly precipitated.

### Pyrophosphate-mediated repair of 2**′**-3**′**-cyclic-phosphate-terminated RNA sequences

When the phosphodiester of the RNA backbone undergoes nucleophilic attack by an adjacent 2′ oxyanion, the RNA strand is cleaved, generating sequences terminated with a 2′-3′-cyclic phosphate (2′-3′-cP); this damage is the most common type of RNA degradation. This cyclization reaction also occurs in self-cleaving ribozymes, many RNases, and base-catalyzed RNA degradation,^42–44^ and is responsible for the relatively low stability of RNA, when compared with DNA or other nucleic acids that lack a phosphate-adjacent hydroxyl group. Because the 2′-3′-cP acts as a chain terminator and prevents the 3′ hydroxyl from participating in extension during polymerization, it represents a replication dead end (Fig. 4a,b). Thus, we tested the polymerase ribozyme for pyrophosphate-mediated damage repair of an RNA hairpin terminated with a 2′-3′-cP. The cyclic phosphate can be removed enzymatically by T4 polynucleotide kinase (PNK), which we used as a positive control for testing the extension efficiencies of the hairpin substrates (Extended Data Fig. 1).^38,39^

**Figure 4.**
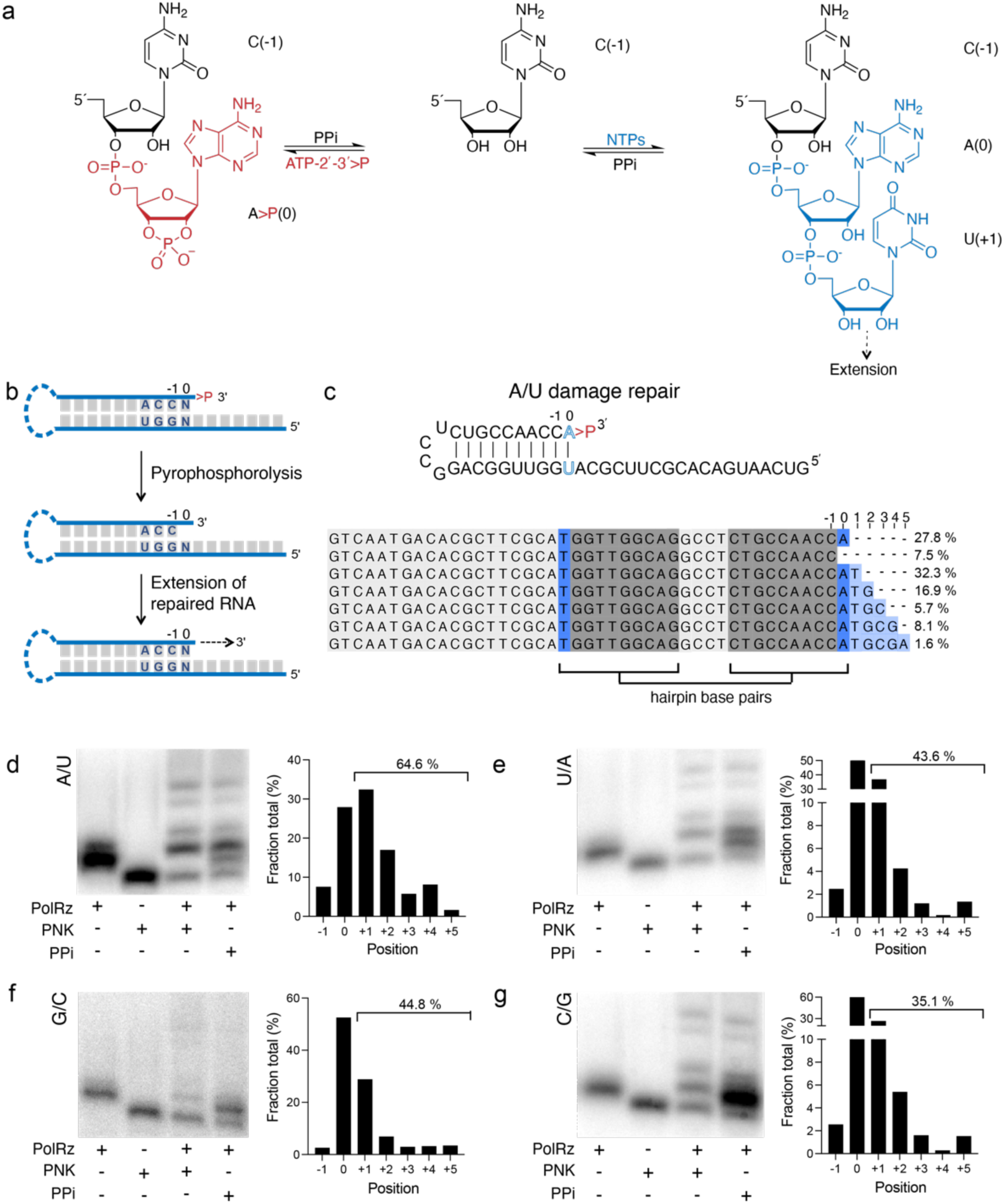
Pyrophosphate-facilitated repair of an RNA damage site by a polymerase ribozyme. (a) A hairpin substrate terminated with a 2′-3′-cyclic phosphate (2′-3′-cP; red nucleotide) cannot be extended by the polymerase ribozyme. However, upon removal of the terminal adenosine by ribozyme-mediated pyrophosphorolysis, the revealed 3′-OH becomes available for further addition of a nucleoside (blue) from a “correct” NTP. (b) Graphical representation of a hairpin with 2′-3′-cP damage and subsequent repair *via* pyrophosphorolysis. (c) Full sequence of an A/U hairpin and aligned reads from high-throughput sequencing analysis showing removal of the terminal adenosine (dark blue), and subsequent extension up to five bases (light blue). Percentages on the right indicate the fraction of each sequence in the dataset. The numbering above the alignment represents terminal nucleotides defined in a. (d-g) Analysis of ^32^P-ATP-labeled 2′-3′-cP-damaged hairpins terminated in all four base-pairs; terminal base-pairs are indicated alongside each PAGE analysis (left-hand image of each set). Graphs on the right of each panel show the percentage of each sequence as determined from HTS data, starting with sequences in which the damaged terminal nucleotide has been removed (fraction of sequences ending at position −1) and proceeding to extensions of the repaired RNA (0–5). The starting RNA containing the 2′-3′-cP cannot be ligated to the primer used for sequencing and is therefore not included in this analysis.

We prepared RNA hairpins terminated with each of the four canonical nucleotides base-paired to the matching nucleotide on the template strand and terminated with a 2′-3′-cP. As expected, no polymerization by the ribozyme was observed when extension of the 2′-3′-cP-damaged sequence was attempted. However, upon addition of pyrophosphate, the terminal damaged nucleotide was removed, and each of the sequences could be extended by the polymerase ribozyme (Fig. 4c-g, Extended Data Fig. 8–9, and Supplementary Fig. 4). The sequences could also be extended after removal of the 2′-3′-cP by PNK. Both extension and repair were largely dependent on the sequence, with the U/A hairpin undergoing repair the most efficiently.

We hypothesized that the mechanism of damage repair proceeds through pyrophosphorolysis of the 3′ terminal 2′-3′-cP-damaged nucleotide, as described earlier in the context of mismatch repair. Therefore, we analyzed the pyrophosphate-mediated removal of the terminal nucleotide by high-resolution PAGE but did not observe a gel mobility change that could be unequivocally attributed to removal of a single nucleotide from the 2′-3′-cP-terminated hairpin (see SI note 1). To better understand pyrophosphate-mediated repair of damaged RNAs, we began by performing repair experiments that could indirectly indicate the removal of the terminal nucleotide. First, we omitted the rNTP that corresponded to the damaged nucleotide in the extension reaction. For example, in the case of an A/U hairpin, we omitted ATP from the rNTPs mixture, because following excision of the terminal AMP-2′-3′-cP, the ribozyme would need to replace it with undamaged AMP to continue the extension reaction. As expected, we found that if ATP was not present in the reaction, the sequence was not extended (Supplementary Fig. 5). To gather additional evidence of damage repair, we transcribed hairpins without ^32^P and performed the polymerization reaction with *α*-^32^P-NTPs to incorporate the ^32^P tracer. We observed that the *α*-^32^P-NTP is only utilized in sequences that have been repaired, and although the ribozyme can incorporate it as a mismatched nucleotide during the extension, this occurs with lower efficiency (Supplementary Fig. 6). Finally, we observed no reaction in the presence of only uridine and pyrophosphate, indicating that a non-canonical reaction, such as opening of the 2′-3′-cP by pyrophosphate to form a triphosphate that can subsequently react with unphosphorylated uridine, did not take place (Supplementary Fig. 7).

### Repair of both cyclic-phosphate damage and mismatches

For certain hairpins, we observed both damage and mismatch repair upon addition of pyrophosphate. Specifically, the purine/purine mismatches (A/A, G/A, and A/G) and the G·U wobble pairs (G/U and U/G) could be repaired by the ribozyme when pyrophosphate was added to the extension reaction of these 2′-3′-cP-terminated mismatches (Extended Data Fig. 10). These reactions exemplify a repair and copying of RNA strands that were mismatched, cleaved, or degraded and formed imperfect primer−template pairs either in *cis* or in *trans*, enabling replication of novel segments or duplication of existing segments as a pathway in genetic innovation.

## Discussion

Modern biological systems possess distinct complexes and pathways dedicated to repairing different types of genome damage that could lead to degradation of the genetic information. Early replicating systems in an RNA world would not have had those functionalities or enzymatic diversity available to them, but damage and mismatch repair would likely have been necessary to replicate long genomes. It has been widely hypothesized that for known polymerase ribozymes to truly achieve efficient self-replication capability, an increase in the current standard of fidelity of replication is necessary. Though early studies on the first polymerase ribozymes hypothesized that pyrophosphorolysis could improve fidelity, this reactivity was not detected.^34^ We address this gap by demonstrating repair activity in the presence of pyrophosphate by an evolved polymerase ribozyme originally selected to extend a primer along a template in an effort to generate a system of self-sustained replication.^18^ We show that the tC19Z polymerase ribozyme can repair mismatched sequences, which presents a possible solution to the fidelity problems faced by early replication ribozymes: in a reaction with a long extension product, we observed an increase in the average length of sequences and an overall higher fidelity in the presence of millimolar pyrophosphate. In addition, tC19Z can use pyrophosphate to repair 2′-3′-cyclic-phosphate-damaged RNA sequences that are formed *via* base-catalyzed RNA cleavage. Although many ribozymes exhibit varying levels of replication efficiency, under saturating pyrophosphate conditions, tC19Z is the first ribozyme with a demonstrated capability of repairing RNA.

Although our findings established that tC19Z can repair terminal mismatched sequences, the extent of ribozyme-mediated repair is highly dependent on mismatch identity, and distinct trends are present. Previously published findings suggested that less-evolved polymerase ribozymes (more closely resembling the class I ligase) also exhibit nucleotide preferences and add mismatches or extend mismatches with varying efficiencies.^15^ Our observations on single nucleotide incorporation and patterns of extended sequences show that tC19Z has preferences for some nucleotides and sequences over others. For example, the ribozyme will add to uridine nucleotides in preference to other nucleotides, and similarly will repair damaged uridine nucleotides with greater frequency. Based on the mismatch repair trends, we hypothesize that there is a structural foundation for the trends in ribozyme-mediated extension of mismatched sequences and the effect of pyrophosphate on extension. Specifically, the width of the terminal base-pair appears to affect the recognition and polymerization capabilities of the ribozyme ^45–49^. Although a recent cryo-EM structure has provided new insight into the structural basis for the function of polymerase ribozymes, additional structural data for this ribozyme-hairpin RNA complex are needed to definitively conclude what the ramifications of mismatches are on the ribozyme’s polymerization reactivity.^50^

The relationship of sequence length and replication fidelity was originally outlined by Eigen, stating that as conserved sequence length increases so must replication fidelity to maintain functionality over successive rounds of copying, and hypothesizing that a mechanism of repair would be critical in an evolving world.^51,52^ Our sequencing analysis reveals a connection between downstream fidelity and mismatch repair: a substantial increase the downstream fidelity of extended sequences is observed when mismatches are corrected. Some mismatches negatively affect the fidelity of downstream polymerization more prominently than others, indicating that if the ribozyme inserts one incorrect base, it is more likely to add additional mismatches. For example, if the U/C mismatched hairpin is not repaired, over 60% of subsequently extended sequences have a second mismatch in the +1 position, and the majority involve addition of a cytosine to make a C/A mismatch. Furthermore, in the long extension experiments, shorter sequences present in the data exhibited a varying number of mismatches at their 3′-end. Ultimately, compounding downstream mismatches leads to termination of replication, which would result in genome replication catastrophe in an evolving world. By improving fidelity through a ribozyme-mediated repair mechanism, we show a new functionality accessible to an evolving system that can directly address the theoretical limits on replicating longer sequences that maintain their function.

Our study establishes that a single ribozyme can perform several distinct biochemical functions, likely with the same active site.^16,19,23^ Such economy of function may be a hallmark of emerging genome-encoded functional macromolecules before dedicated synthesis and repair machinery evolved. These results bring us one step closer to generating truly self-replicating systems that could have evolved in an RNA world.

## Materials and Methods

### Cloning tC19Z into a pUC19 vector

The insert, tC19Z, was synthesized and obtained from Integrated DNA Technologies and then was subsequently amplified with primers containing the restriction enzyme sites BamHI and EcoRI (all enzymes were obtained from New England BioLabs) (**Supplementary Table 1**; sequence #1 is the ribozyme construct).^18^ The tC19Z insert and pUC19 vector were double digested with BamHI and EcoRI restriction enzymes for 4 hours. Additionally, the pUC19 vector was further treated with calf-intestinal, alkaline phosphatase (CIP) for 1 hour following the double digestion. The insert was ligated into the vector using T4 DNA ligase and incubating at 16 °C for 16 hours. The resulting ligation was transformed into DH5*α* competent cells and subsequently plated on ampicillin-containing agar plates which were grown at 37 °C for 14 hours. Single colonies were picked and grown in 3 mL of ampicillin containing LB broth for 14 hours. Cultures were subjected to alkaline lysis and the plasmids were purified. The presence of tC19Z in the vector was confirmed by Sanger sequencing. Future polymerase chain reactions (PCR) of tC19Z were performed fresh from the plasmid for each batch of ribozyme.

### PCR and transcription of tC19Z and generation of hairpin constructs

PCR was performed to isolate tC19Z from the plasmid prepared above. The plasmid (5 ng), each primer (0.5 µM) (**Supplementary Table 1**; primers # 4 and 5), and *Taq* DNA polymerase were used to amplify the sequence at an annealing temperature of 49 °C. PCR products of tC19Z from the plasmid prepared above were used for *in vitro* transcription. Transcription was performed under the following conditions: 50 mM Tris HCl, 2 mM spermidine, 12 mM MgCl_2_, 8 mM DTT, 2 mM each rNTP, and 0.01% Triton X-100 in a total volume of 100 µL. T7 RNA polymerase was added to the reaction and allowed to react at 37 °C for 3 hours. The transcriptions were desalted with G15 Sephadex and purified using PAGE [18% w/v acrylamide/bisacrylamide 19:1 solution, 7 M urea, and 0.5x TBE Buffer (50 mM tris(hydroxymethyl)aminomethane hydrochloride, 45 mM boric acid, and 0.5 mM EDTA, pH 8.4)]. RNA was detected by UV shadowing and then was (i) eluted in 300 µL of 300 mM KCl from gel pieces, (ii) precipitated by adding 700 µL of ethanol, and (iii) incubated at –20 °C overnight. The samples were then centrifuged to pellet the precipitate, which was subsequently dried in air and resuspended in water.

To prepare the RNA hairpin substrates for the tC19Z polymerase ribozyme (**Supplementary Table 1**), the hairpin sequences (0.5 µM primers #7–#22; **Supplementary Table 1**) were generated as fusions with the drz-Mbac-1 HDV-like ribozyme (0.5 µM; #6 construct; **Supplementary Table 1**) using primer extension.^37^ Following amplification, the sequence were transcribed in the presence of α-^32^P[ATP] (PerkinElmer) by adding 50 mM Tris HCl, 2 mM spermidine, 12 mM MgCl_2_, 8 mM DTT, 2 mM GTP, 2 mM CTP, 2 mM UTP, 200 µM ATP, and 0.01% Triton X-100 to a 50 µL reaction containing T7 RNA polymerase, and the mixture was allowed to react at 37 °C for 2 hours. The HDV-like ribozyme self-cleaves co-transcriptionally to yield cyclic-phosphate-terminated hairpins. The transcriptions were desalted with Sephadex G10 spin columns (10 mM Tris, 1 mM EDTA, pH 7.2) purified using PAGE, and worked up as described above to generate internally ^32^P-labeled polymerase substrates.

### Adding ^32^P-phosphate to an RNA primer

An RNA primer 10 nucleotides in length was obtained from Dharmacon (primer #2; **Supplementary Table 1**). A ^32^P radiolabel was added to the 5′-end of the sequence by adding the RNA primer at a concentration of 5 µM to a solution containing γ-^32^P[ATP], 1x T4 polynucleotide kinase buffer, and 5 units T4 polynucleotide kinase. The reaction was incubated at 37 °C for 2 hours, purified using PAGE, and worked up as described above.

### Kinetic analysis of pyrophosphate inhibition of RNA primer extension

Kinetic data for pyrophosphate inhibition were obtained by monitoring the extension of a ^32^P-labeled primer by tC19Z over time and varying concentrations of pyrophosphate. Equimolar concentrations (0.5 µM) of tC19Z, primer, and template were mixed and heated to 82 °C for 2 minutes, then cooled slowly to 17 °C for 10 minutes. Varying concentrations of pyrophosphate were then added to the reactions followed by buffer containing 200 mM MgCl_2_, 50 mM Tris HCl buffer (pH 8.3), and 4 mM rNTPs, and the solution was then allowed to incubate at 17 °C. Aliquots (5 µL) were removed at various time points and were quenched with 250 mM EDTA and 7 M urea gel-loading buffer. The reactions were analyzed using an 18% denaturing PAGE to obtain single-nucleotide-resolution images. The gels were imaged on a Typhoon 9410 Variable Mode Imager, and the pixel density of the bands was analyzed using ImageJ to generate the kinetic data. Initial rates were calculated using the first four time points and used to demonstrate pyrophosphate inhibition.

### Repair of cyclic phosphate damaged RNA hairpins

Hairpin substrate repair was visualized and quantified by imaging the extension of an internally ^32^P-labeled hairpin by tC19Z. For negative controls, equimolar concentrations (0.5 µM) of tC19Z and 2′-3′**-**cyclic-phosphate-terminated hairpins were mixed and heated to 82 °C for 2 minutes, and then cooled slowly to 17 °C for 10 minutes. At this point, one of two buffers was added (1^st^ buffer: 200 mM MgCl_2_ and 50 mM Tris buffer; 2^nd^ buffer: 200 mM MgCl_2_, 50 mM Tris buffer, and 4 mM rNTPs), and the reactions were incubated at 17 °C for 24 hours. The reactions were quenched with 250 mM EDTA and 7 M urea gel-loading buffer, and then were analyzed on a 15% PAGE gel. For the repair reaction, equimolar concentrations (0.5 µM) of tC19Z and cyclic-phosphate-terminated hairpin were mixed and heated to 82 °C for 2 minutes, followed by slow cooling to 17 °C for 10 minutes and subsequent addition of 10 mM pyrophosphate. A buffer containing 200 mM MgCl_2_, 50 mM Tris buffer, and 4 mM rNTPs was added, and the reaction was allowed to incubate at 17 °C for 24 hours. The reactions were quenched with 250 mM EDTA and 7 M urea gel-loading buffer and were analyzed on a 15% denaturing polyacrylamide gel. To generate a positive control sample, the cyclic phosphate was removed from the hairpin by treating with polynucleotide kinase in 1x PNK buffer (NEB) for 1 hour at 37 °C then inactivated at 65 °C for 10 minutes. The PNK-repaired hairpin substrate was then added to the tC19Z polymerase ribozyme, both at concentrations of 0.5 µM, and the solution was heated to 82 °C for 2 minutes and then cooled slowly to 17 °C for 10 minutes. Next, one of two buffers was added (1^st^ buffer: 200 mM MgCl_2_ and 50 mM Tris buffer; 2^nd^ buffer: 200 mM MgCl_2_, 50 mM Tris buffer, and 4 mM rNTPs), and the reactions were allowed to incubate at 17 °C for 24 hours. The reactions were quenched with 250 mM EDTA and 7 M urea gel-loading buffer and were analyzed on a 15% denaturing polyacrylamide gel. The gels were imaged on a Typhoon 9410 Variable Mode Imager.

### Kinetic analysis of cyclic-phosphate damage repair

The kinetics of tC19Z-mediated extension of pyrophosphate-repaired hairpin, PNK-repaired hairpin, and pyrophosphate inhibited PNK-repaired hairpin were obtained under similar conditions as those described above. Either hairpin or PNK-repaired hairpin was incubated with tC19Z at 82 °C for 2 minutes and then cooled slowly to 17 °C for 10 minutes. For the reactions containing pyrophosphate-repaired hairpin and pyrophosphate-inhibited hairpin, pyrophosphate was added at a concentration of 10 mM. A buffer containing 200 mM MgCl_2_, 50 mM Tris buffer, and 4 mM rNTPs was added to each reaction, and the solutions were allowed to incubate at 17 °C. Time points were removed from the reactions in 5 µL aliquots and quenched with 250 mM EDTA and 7 M urea gel-loading buffer. The reactions were analyzed on a 15% denaturing polyacrylamide gel that was imaged on a Typhoon 9410 Variable Mode Imager. ImageJ was used to quantify band density and generate the kinetic data for tC19Z extension of hairpins under varying pyrophosphate and repair conditions.

### Hairpin migration and extension under various conditions

To better understand hairpin migration and reactivity, an A/U hairpin was either left untreated or allowed to react with an alkaline phosphatase [5 µM hairpin, 1x Cutsmart buffer (NEB), and shrimp alkaline phosphatase (NEB)] for 30 mins at 37 °C. The two conditions were then either left untreated, treated with PNK as described above, or incubated with acid (25 mM HCl) followed by quenching. Each solution was then analyzed on a 15% denaturing polyacrylamide gel. Furthermore, each treated hairpin (0.5 µM) was heated with tC19Z (0.5 µM) for 2 minutes at 85 °C and then cooled to 17 °C over 10 minutes. A buffer containing 200 mM MgCl_2_ and 50 mM Tris HCl (and, in some reactions, 4 mM rNTPs) was added. Additionally, 4 mM uridine and/or 10 mM pyrophosphate were added to some reactions. The mixtures were incubated at 17 °C for 24 hours followed by quenching with 250 mM EDTA and 7 M urea gel-loading buffer. The reactions were analyzed on a 15% denaturing polyacrylamide gel that was imaged on a Typhoon 9410 Variable Mode Imager.

### Incorporation of ^32^P-rNMPs into non-labeled, repaired hairpins

To further demonstrate repair, an A/U hairpin was transcribed with an α-^32^P[ATP] or without (cold). Some of the hairpins were then treated with PNK as described above. The ^32^P-labeled hairpin (0.5 µM) was incubated with tC19Z (0.5 µM) at 82 °C for 2 mins and cooled to 17 °C over 10 minutes. A buffer containing 200 mM MgCl_2_ and 50 mM Tris HCl (and, in some reactions, 4 mM rNTPs) was added. To some reactions was added either 10 mM pyrophosphate or α-^32^P[ATP] and 10 mM pyrophosphate. The cold hairpin was treated in the same way. The mixtures were incubated at 17 °C for 24 hours followed by quenching with 250 mM EDTA and 7 M urea gel-loading buffer. The reactions were analyzed on a 15% denaturing polyacrylamide gel that was imaged on a Typhoon 9410 Variable Mode Imager.

A U/A hairpin substrate was transcribed without α-^32^P-ATP and either left untreated or allowed to react with PNK as described above. The hairpin (0.5 µM) was heated with tC19Z (0.5 µM) for 2 minutes at 85 °C and then cooled to 17 °C over 10 minutes. A buffer containing 200 mM MgCl_2_, 50 mM Tris HCl and, in some reactions, 10 mM pyrophosphate was added. Additionally, either α −^32^P[ATP], α −^32^P[UTP], or α −^32^P[CTP] was added. The reactions were incubated at 17 °C for 24 hours followed by quenching with 250 mM EDTA and 7 M urea gel-loading buffer. The reactions were analyzed on a 15% denaturing polyacrylamide gel that was imaged on a Typhoon 9410 Variable Mode Imager.

### Extension of mismatched RNA hairpin sequences

Mismatched RNA hairpins were extended by the tC19Z polymerase ribozyme under conditions of PNK repair or pyrophosphate. Pyrophosphate facilitated extension of both a cyclic-phosphate-damaged hairpin and a mismatched hairpin when 0.5 µM hairpin was added to 0.5 µM tC19Z. The solution was incubated at 82 °C for 2 minutes and then cooled slowly to 17 °C for 10 minutes. Pyrophosphate was added at a final concentration of 10 mM, followed by a buffer containing 200 mM MgCl_2_, 50 mM Tris buffer, and 4 mM rNTPs, and the solution was incubated at 17 °C for 24 hours. A reaction containing a PNK-repaired mismatched hairpin and tC19Z at concentrations of 0.5 µM each was incubated at 82 °C for 2 minutes and then cooled slowly to 17 °C for 10 minutes. Pyrophosphate was added at a concentration of 10 mM, followed by a buffer containing 200 mM MgCl_2_, 50 mM Tris buffer, and 4 mM rNTPs, and the reaction was allowed to proceed at 17 °C for 24 hours. All reactions were quenched with 250 mM EDTA and 7 M urea gel-loading buffer and then analyzed on a 15% denaturing polyacrylamide gel that was imaged on a Typhoon 9410 Variable Mode Imager. ImageJ was used to quantify band density to generate fraction extended graphs.

### High-throughput sequencing of RNA hairpins

RNA hairpins were analyzed by Illumina Sequencing to determine sequence identity. Cyclic phosphate repair and mismatch extension reactions, as described above, were performed and fractionated by PAGE. The images obtained were analyzed for extension products, and the corresponding bands were excised from the gel, eluted, and precipitated with ethanol. The RNA sequences were resuspended in water and then ligated to a DNA sequence containing an Illumina forward adapter (#24 and 25, **Supplementary Table 1**). The DNA adapter sequence was designed with a 5′-phosphate, followed by four random nucleotides to reduce ligation bias at the 3′-RNA terminus. The adapter also contained a 3′-amine to prevent self-ligation. The oligo (10 µM) was added to a mixture containing PEG 8000, 1x T4 RNA ligase buffer, 15 µM ATP, and T4 RNA ligase 1 to a total volume of 100 µL to adenylylate the 5′-end of the DNA sequence (5′-App oligo). G15 Sephadex was used to desalt and remove residual ATP from the 5′-App oligo.

To anneal the sequences, the 5′-App oligo was added in approximately three times excess of the RNA hairpin, and the resulting solution was heated to 95 °C for 3 mins and then cooled to 25 °C at a rate of 1 °C/min. To ligate, the annealed oligos were added to 1x T4 RNA ligase buffer, PEG 8000, and T4 RNA ligase 2, truncated KQ to a total volume of 20 µL, and allowed to react at 25 °C for 18 hours and then at 4 °C for 2 hours.

To a 5 µL aliquot of the ligation reaction was added 2.5 µM of the RT primer (#29 in **Supplementary Table 1**), which was annealed at 75 °C for 3 mins, 37 °C for 10 mins, and 25 °C for 10 mins. Following annealing, 1x ProtoScript II buffer, 0.5 mM each dNTP, 5 mM MgCl_2_, 10 mM DTT, and ProtoScript II were added and the reverse transcription reaction was heated to 50 °C for 2 hours to generate cDNA.

The Illumina forward adapter was added to the cDNA through a PCR reaction (using primer #31 in **Supplementary Table 1**). Forward and reverse primers (#31 and #30; 1 µM) were added to the cDNA, as well as 1x Q5 reaction buffer, 5 mM of each dNTP, and Q5 DNA polymerase. The reaction was annealed at 52 °C and extended at 72 °C. The Illumina barcodes were added in a second PCR reaction containing 1 µM of each primer, 1x Q5 buffer, 5 mM each dNTP, and Q5 DNA polymerase. The reaction to install the Illumina barcodes was annealed at 65 °C and extended at 72 °C, and the reaction was purified via agarose gel. The libraries were submitted for Illumina Sequencing. PhiX was removed from the resulting reads using bowtie2, and the remaining reads were trimmed using CutAdapt to remove adapter regions. The 5′-adapter (TCTACACTCTTTCCCTACACGACGCTCTTCCGATCT) was removed first, then the 3′-adapter (GATCGGAAGAGCACACGTCTGAACTCCAGTCAC), with at most 10% error in single-end mode. Sequences were counted in FastAptamer using fastaptamer_count. Starting and extended sequences were extracted from data for analysis in FastAptamer using fastaptamer_search for each sequence with unique identity and length.^53^ Specifically, the two-nucleotide tag (AC or GT) incorporated from sequences #24 or #25, followed by four random N nucleotides, and ending with the sequence of interest were searched for and compiled. These sequences were then curated visually in Jalview. Sequence identity was analyzed at each position and the number of sequences in each case were used to calculate extension efficiency, pyrophosphorolysis, and fidelity (SI Note 2). The sequencing was performed on one representative reaction for each condition, as is convention in the field when determining polymerase ribozyme fidelity.

### Long-read primer extension and sequencing

For sequencing the long-read experiments, we prepared an RNA template corresponding to a sequence previously used with the tC19Z rRNA polymerase ribozyme.^18^ The sequence was transcribed from the long-read DNA template (sequence #23 in **Supplementary Table 1**) prepared as a double-stranded DNA using the template strand and a T7 RNAP primer-extended as described above, and PAGE-purified, as described above. The template RNA was combined with the RNA primer (sequence #2 in **Supplementary Table 1**) and the tC19Z ribozyme at 1 µM concentration (each) and this mixture was incubated at various concentrations of pyrophosphate for 4 days at 17 °C as described above. The reactions were desalted using Sephadex G10. The desalted samples were not further purified. The samples were then ligated to the ligation primer for sequencing (sequences #24 and #25 in **Supplementary Table 1**) and reverse-transcribed as described above. The cDNAs were then ligated to the cDNA ligation hairpin (sequence #26 in **Supplementary Table 1**), which contains a 7-nt random sequence to allow splint-ligation by a DNA ligase independent of the 3′ terminal sequence of the cDNA and allow amplification of all RNAs present in the samples, as described previously.^54^ To suppress self-ligation, the cDNA ligation hairpin DNAs contained 3′ terminal carbon spacer. The samples were incubated in T4 DNA ligase buffer (NEB) with 1 mM ATP, 0.5 µL of T4 DNA ligase, 0.5 M betaine, 20% PEG 8000, and 0.5 µL of RNaseH (NEB). To preferentially amplify the ribozyme-extended RNA primers, a DNA primer that overlapped with the RNA primer sequence (CUGCCAACCG) extended by the ligated hairpin sequence was designed and used for PCR amplification (sequence #27 in **Supplementary Table 1**) together with the RT primer for sequencing (#29 in **Supplementary Table 1**). A nested PCR reaction was performed to further enrich for sequences of the RNA primer–template ribozyme extension reaction and introduced the Illumina reverse primer sequence (#28 in **Supplementary Table 1**) and the reactions were sequenced as described above. Whereas reactions were designed to bias towards amplification of the RNA primer-extensions, we observed many other amplicons, including many sequences that corresponded to products of primer or hairpin self-ligations.

The libraries were sequenced by Illumina Sequencing, and the resulting reads were trimmed using CutAdapt to remove primer regions. Sequences were counted in FastAptamer using fastaptamer_count. Sequences containing the primer sequence were extracted from the data for analysis in FastAptamer using fastaptamer_search for each sequence and subsequently visually aligned and curated using Jalview. Sequence lengths and fidelity were obtained using Jalview (SI Note 2). The sequencing was performed on one representative reaction for each condition, as is convention in the field when determining polymerase ribozyme fidelity.

### Measuring ATP concentrations using a firefly luciferase assay

A solution containing a 2-µM PNK-repaired A/C hairpin and 0.5-µM tC19Z polymerase ribozyme was incubated at 82 °C for 2 minutes and then cooled slowly to 17 °C for 10 minutes. Pyrophosphate was added at a concentration of 10 mM followed by a buffer containing 200 mM MgCl_2_ and 50 mM Tris buffer, and the reaction was allowed to proceed at 17 °C for 3 days. Control reactions were incubated as above, but in the absence of individual components, as indicated. After 3 days, the reactions were centrifuged, and 0.5 µL of the supernatant was added to 4.5 µL of a solution containing inorganic pyrophosphatase and firefly luciferase (preincubated for 3 h in presence of D-luciferin to deplete any ATP co-purified with the enzyme). Luminescence was generated using the StayBrite Highly Stable ATP Bioluminescence Assay Kit (BioVision), as described previously.^40^ The reaction contained 50 µM D-luciferin, 1 µL of 4x StayBrite reaction buffer, and 1 µL StayBrite luciferase enzyme, in addition to 0.5 µL of inorganic pyrophosphatase (NEB). Each reaction was loaded in a well of a black 384-well plate and imaged using a high numerical aperture lens (Voigtländer 29 mm F/0.8) mounted on an EMCCD camera (iXon Ultra 888, Andor). The temperature of the camera electronics was maintained at –100 °C during detection using a thermoelectric recirculating chiller (Thermocube 300, Solid State Cooling Systems). During imaging, 10 one-minute exposures were collected. The images were then stacked, a Z-projection was generated, and relative luminescence was measured using ImageJ. To generate a standard curve, different ATP concentrations in nM range were incubated in the same buffer system, including pyrophosphate. For the no-pyrophosphate control, pyrophosphate was added to the samples at the end of the experiment and the samples were treated so that the same amount of soluble pyrophosphate was present in each sample when incubated with the pyrophosphatase and luciferase enzymes.

## Acknowledgements

We thank the staff at the Genomics High-Throughput Facility (GHTF) Shared Resource of the Cancer Center, Support Grant (P30CA-062203) at the University of California, Irvine and NIH shared instrumentation grants 1S10RR025496-01, 1S10OD010794-01, and 1S10OD021718-01; and S. Flynn for assistance with sequencing data. We thank C. Langeberg and D. Treco for comments on the manuscript, and J. H. D. Cate for his support. This work was supported by the John Templeton Foundation through a grant to the Foundation for Applied Molecular Evolution (to A. L.), NSF CBET Biophotonics 1804220 (to A. L.), NASA Exobiology 80NSSC21K0488 (to A. L.), NASA ICAR 80NSSC21K0596 (to A.L.), NSF GRFP (to A. K.), Goldwater Scholarship (to L. Y.), and UCI Undergraduate Research Opportunities Program awards (to L. Y. and I. L.).

## Current Affiliations

*California Institute for Quantitative Biosciences, University of California at Berkeley, Berkeley, CA, USA*

Alexandra D. Kent

*Department of Chemistry, Harvard University; Cambridge, MA, USA*

Lucy L. Yang

*Amgen; Thousand Oaks, CA, USA*

Kyle H. Cole

## Author contributions

A.D.K. and A.L. conceived and designed the study. A.D.K. performed the experiments with assistance from L.L.Y., K.H.C. and A. L. The data were analyzed by A.D.K. and A.L. The manuscript was prepared by A.D.K and A.L.

## Data and materials availability

All data are available in the main text, the supplementary materials, or deposited in a sequence repository.

## Extended Data Figures

**Extended Data Fig. 1.**
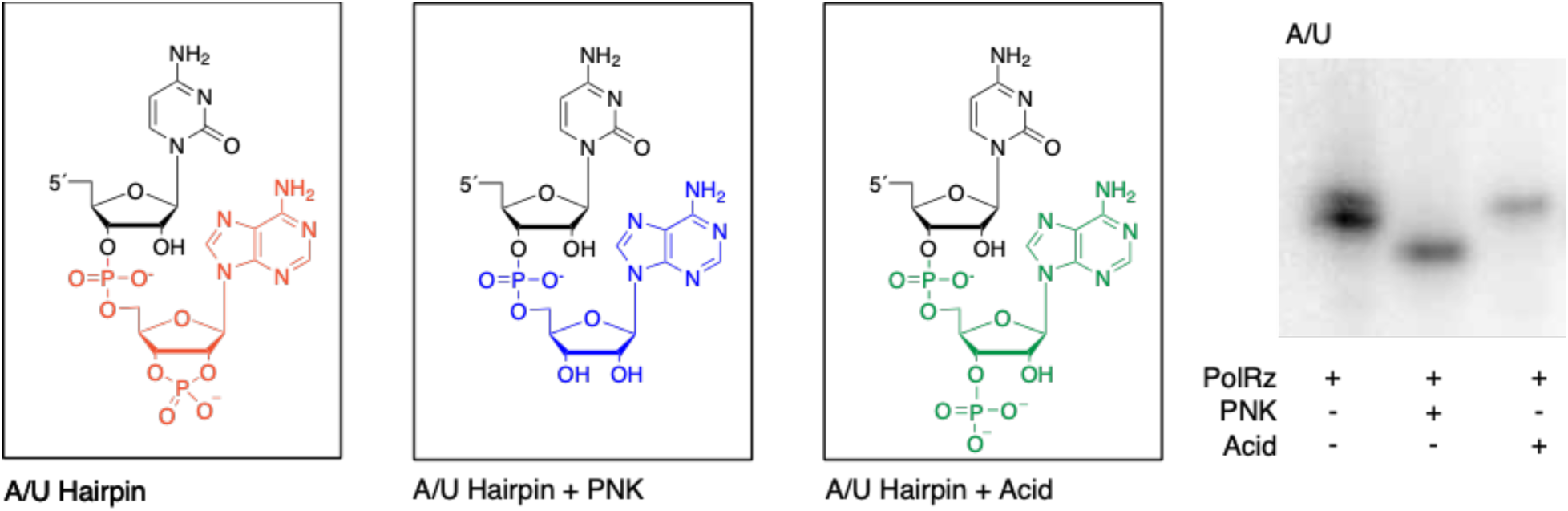
Gel mobility of A/U hairpin. An A/U hairpin has a 2′-3′-cyclic phosphate resulting from drz-Mbac-1 ribozyme self-scission (red and left lane on PAGE gel). After repair with PNK, the cyclic phosphate is removed (blue), and the corresponding band migrates faster on the PAGE gel (middle lane). When the 2′-3′-cyclic-phosphate-terminated hairpin is treated with hydrochloric acid, the cyclic phosphate is partially hydrolyzed, yielding a 3′ monophosphate (green) or 2′ monophosphate (not shown), and the mobility of the corresponding PAGE band is retarded (right lane).

**Extended Data Fig. 2.**
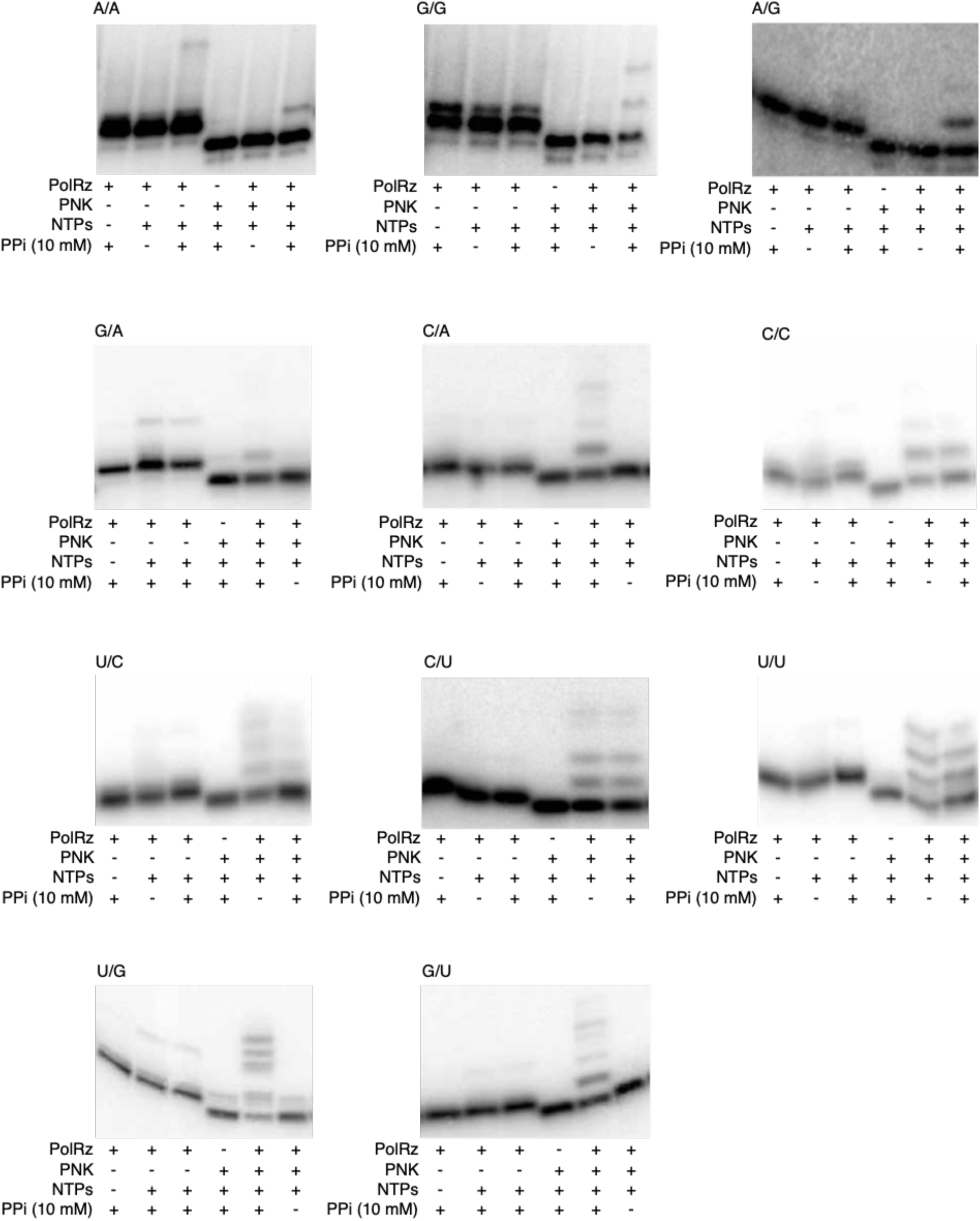
Ribozyme-catalyzed, pyrophosphate-facilitated repair reactions for all mismatch combinations. All mismatches were ^32^P-labeled and reacted under standard conditions with variations indicated with + and − symbols. The reactions were then analyzed by high-resolution PAGE.

**Extended Data Fig. 3.**
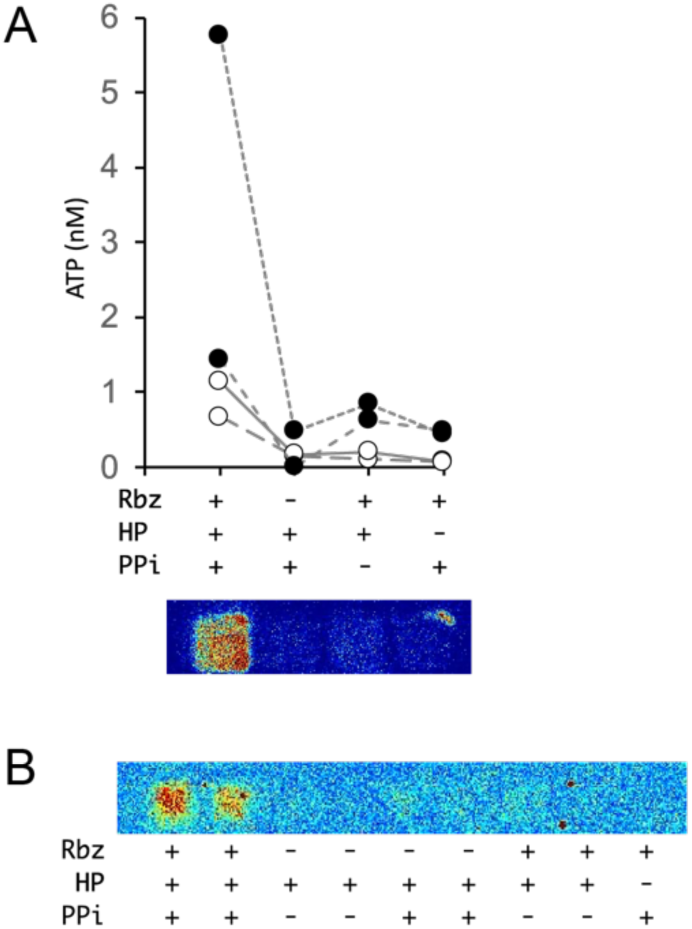
Luciferase-based assay measures ATP produced during ribozyme-catalyzed pyrophosphorolysis. (A) Firefly luciferase assay measuring the concentration of ATP in nM generated by the removal of the terminal adenosine monophosphate of an A/C mismatched hairpin, as calculated by use of a standard curve. Reactions lacking the polymerase ribozyme, hairpin, or pyrophosphate show only background levels of luminescence. Higher concentrations of ATP are present in reactions containing all three components. Open and filled circles represent two independent preparations of the polymerase ribozyme and the hairpin substrate, with filled circles showing higher background ATP levels apparently co-purified with the polymerase ribozyme. Lines connect individual experiments performed in parallel on the same day and from the same stocks. A representative luminescence image is shown below the graph. (B) Additional luminescence image used to generate the values shown in A and showing additional control reactions lacking both the ribozyme and pyrophosphate.

**Extended Data Fig. 4.**
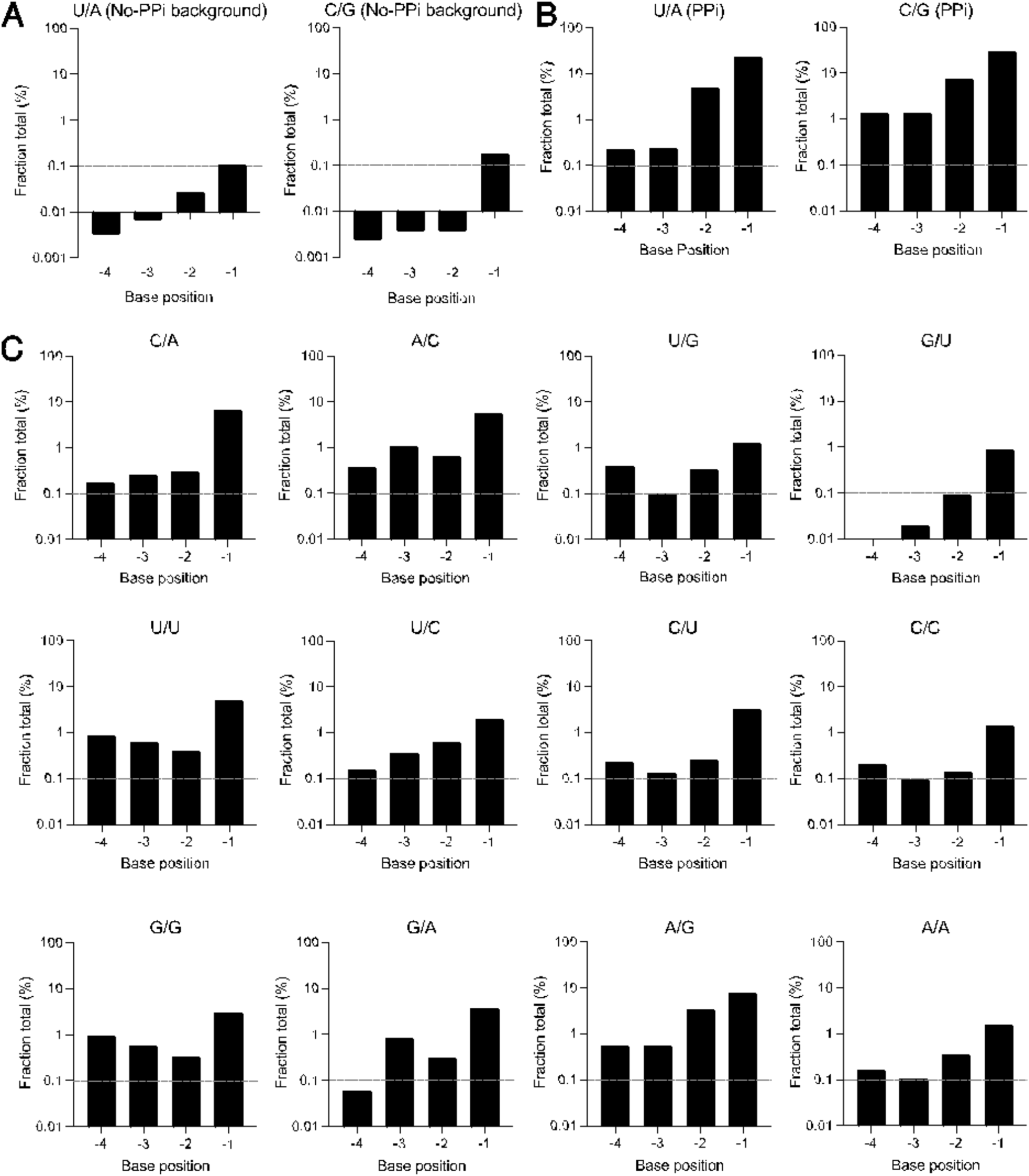
Removal of terminal nucleotides in the presence and absence of pyrophosphate. Addition of pyrophosphate increases the fraction of shorter sequences as determined by high-throughput sequencing. (A) Matched U/A and C/G hairpins terminated with a 2′-3′-cyclic phosphate and incubated in absence of pyrophosphate show very low fractions of shorter sequences, indicating low background of these sequences in the samples. (B) Matched hairpins terminated with a 2′-3′-cyclic phosphate yield a fraction of shorter sequences that is about 2 orders of magnitude higher, particularly in the –1 position, after treatment with ribozyme and pyrophosphate. These results provide direct evidence for ribozyme-catalyzed pyrophosphorolysis of RNA. (C) All mismatched sequences following incubation with pyrophosphate and ribozyme. Some sequences exhibit higher percentages of pyrophosphorolysis than others, largely following the trends described in Figure 4. To ease comparison, all graphs are plotted on the same scale, with the horizontal axis placed at 0.01% and a gray dashed line placed at 0.1%.

**Extended Data Fig. 5.**
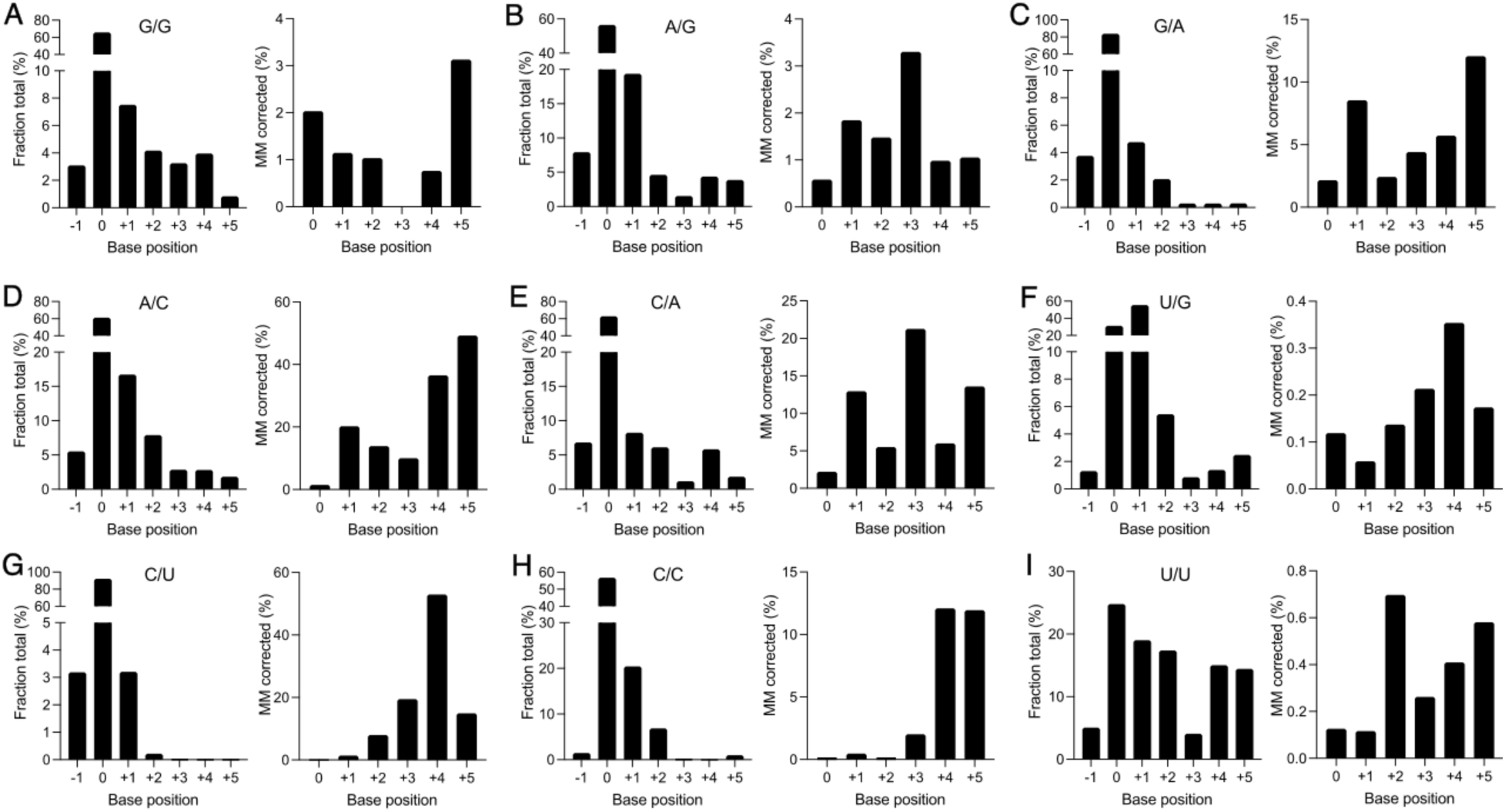
Extension and repair of mismatches. Mismatched hairpins repaired and extended with distinct efficiencies, as determined by high-throughput sequencing. The left graph of each panel shows the fraction of sequences terminated at each position along the template relative to the starting length (0) after incubation with the polymerase ribozyme, pyrophosphate, and rNTPs. All hairpins were treated with PNK to remove the 2′-3′-cyclic phosphate. “–1” corresponds to the fraction of sequences terminated at nucleotides corresponding to the products of pyrophosphorolysis of the mismatch-terminated hairpin. Hairpins terminated in A/G (B), A/C (D), C/A (E), and U/U (I) show particularly high fractions of sequences with the terminal mismatches removed. The right graph of each panel shows the percentage of each extended segment with the starting mismatch repaired. (A-C) Purine/purine mismatches are repaired and extended at low efficiency, but longer products contain high fraction of repaired sequences. (D–F) Purine/pyrimidine mismatches are extended well and have high percentages of repair, except the U/G wobble pair. For example, a hairpin that started with an A/C mismatch (D), showed more than 15% of sequences extended by one nucleotide (+1, left graph) and ∼20 % of these 1-nt extensions had been repaired (+1, right graph). (G–I) Pyrimidine/pyrimidine mismatches are extended with high efficiencies but are not repaired well.

**Extended Data Fig. 6.**
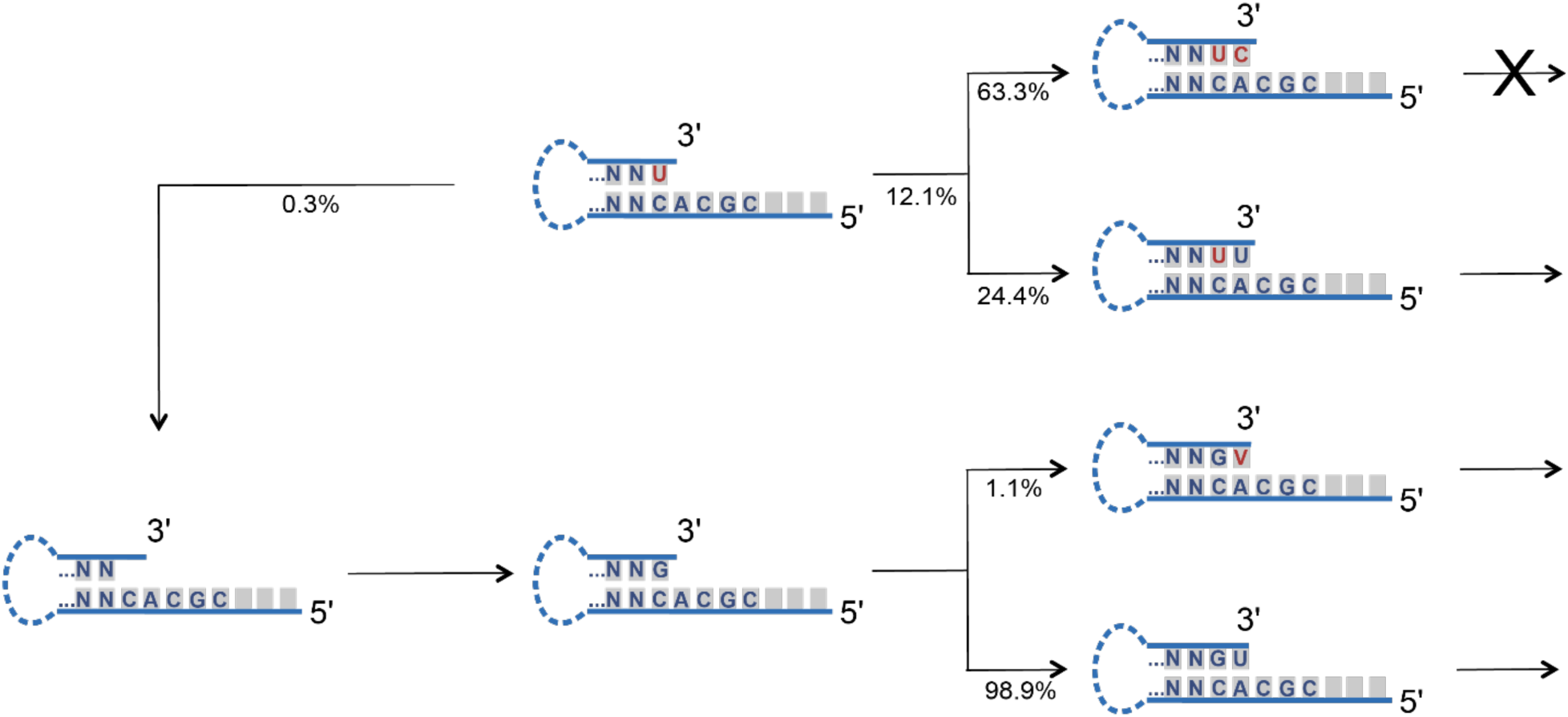
One mismatch can induce addition of more mutations. When a hairpin having a terminal U/C mismatch is incubated with a polymerase ribozyme and tC19Z it can either undergo pyrophosphorolysis or be extended by one nucleotide. In only a low percentage is the mismatch removed, but if it is, the subsequent extension proceeds with high fidelity. If the mismatch is not corrected, the ribozyme will add a second mismatch preferentially which results in chain termination.

**Extended Data Fig. 7.**
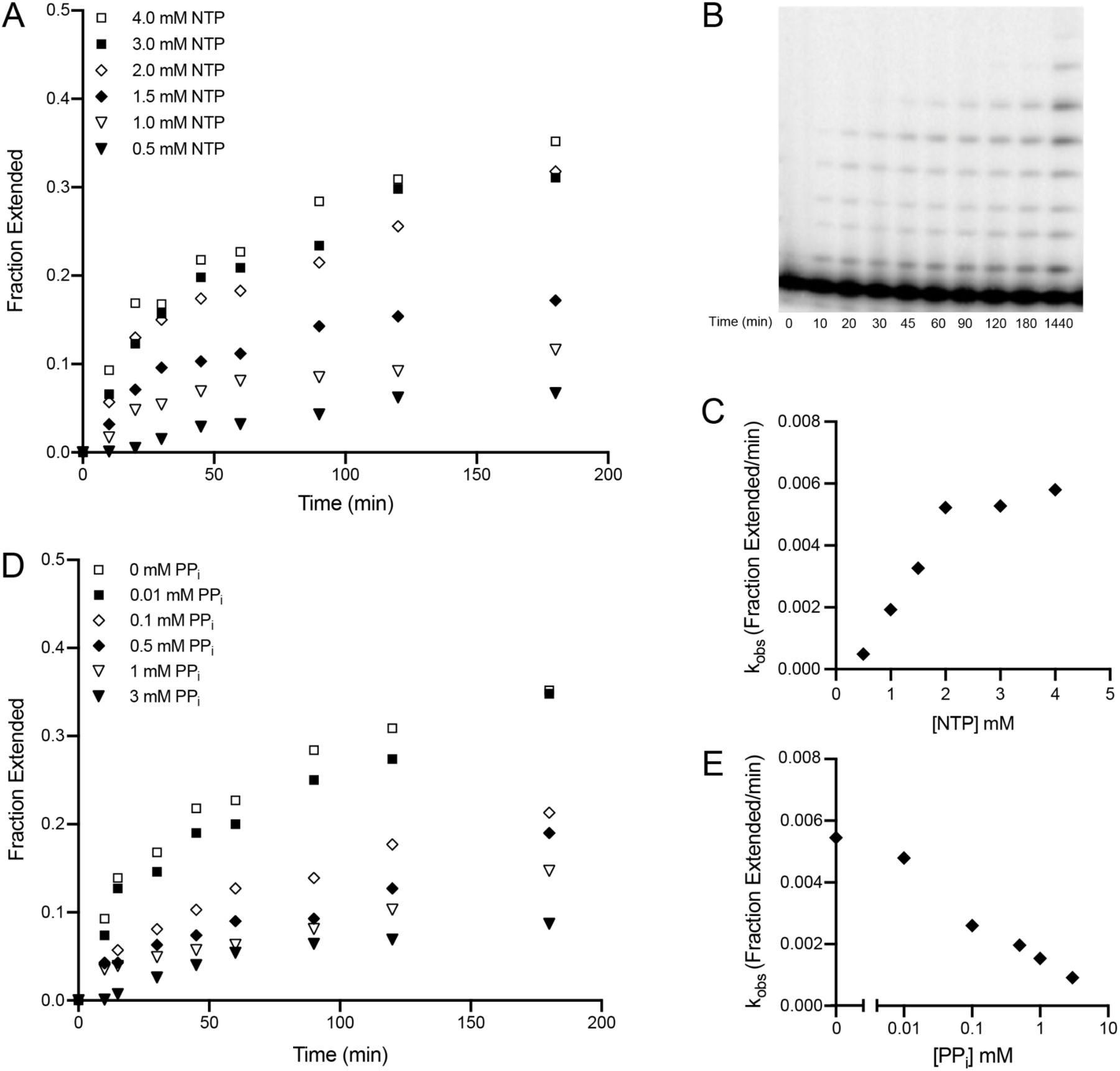
Reaction kinetics of primer extension and pyrophosphate inhibition. (A) The fraction extended of the ^32^P-labeled primer is quantified as a function of time at varying NTP concentrations by high resolution PAGE. (B) High-resolution PAGE of the primer extension at 4 mM rNTPs which is plotted on the graph in A. (C) The fraction of primer extended per minute is plotted as a function of NTP concentration and shows an estimated K_M_ of ∼4 mM. (D) The fraction of the ^32^P-labeled primer extended quantified as a function of time at varying pyrophosphate conditions. (E) The fraction extended per minute is plotted as a function of the log of pyrophosphate concentration.

**Extended Data Fig. 8.**
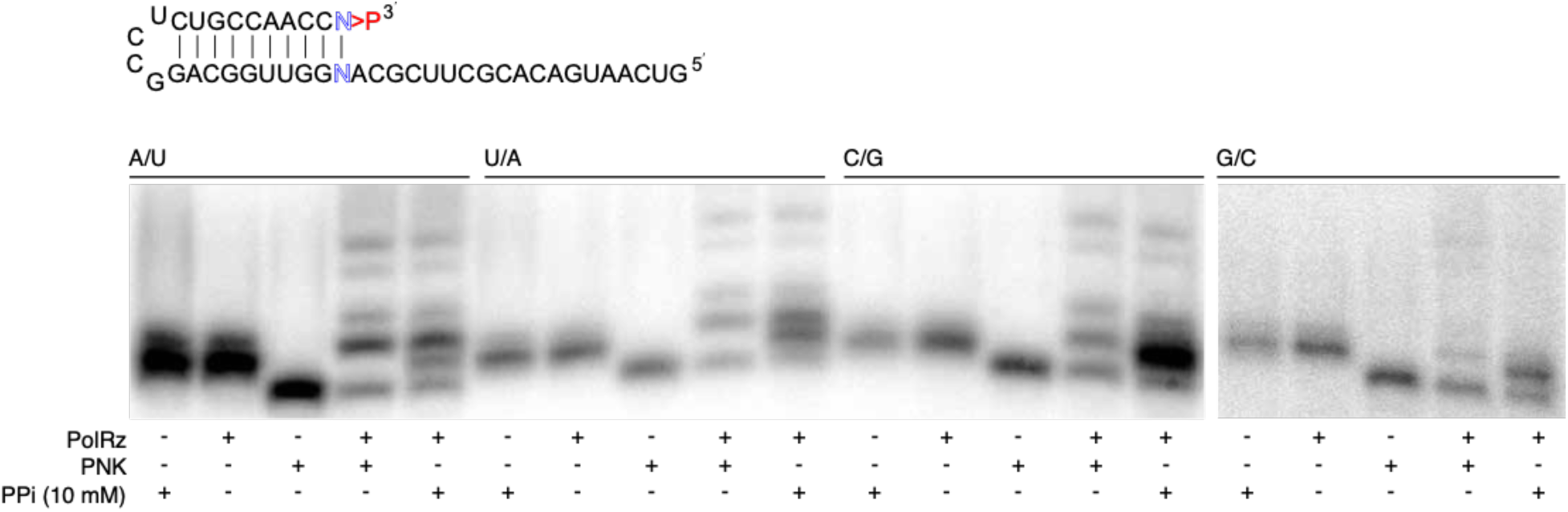
Damage repair reactions for all matched hairpins. The general sequence of the hairpins is shown above the PAGE gels, with terminal base-pair shown as N-N in blue and specific terminal base-pairs indicated above each set of experiments. The site of RNA damage, corresponding to the 2′-3′-cyclic phosphate is indicated as >P in red. All hairpins were allowed to react under standard conditions with reaction conditions variations shown by the + and − symbols. Hairpins were analyzed by high-resolution PAGE. The A/U, U/A, and C/G hairpins were all resolved on the same gel, and the G/C hairpin was resolved on a separate PAGE gel. PNK indicates removal of the terminal 2′-3′-cyclic phosphate by the polynucleotide kinase enzyme to yield canonical 2′ and 3′ hydroxyls, which results in faster migration. All hairpins were extendable by the tC19Z polymerase ribozyme when either treated by the PNK enzyme or pyrophosphate (right two lanes of each set of experiments).

**Extended Data Fig. 9.**
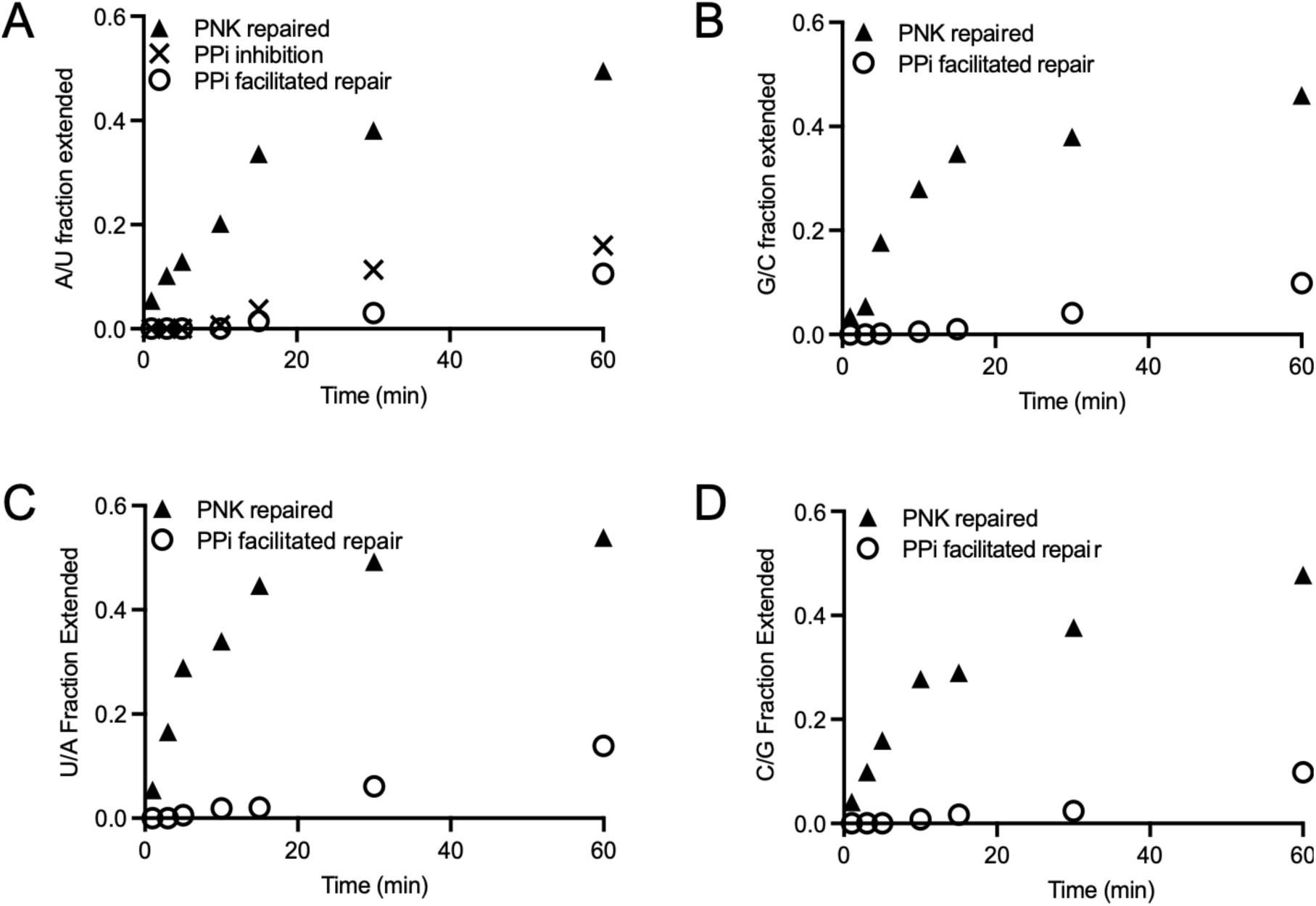
Kinetics of RNA polymerization of damage-repaired substrates. A ^32^P-labeled hairpin terminated with 2′-3′-cyclic phosphate was either treated with PNK or pyrophosphate and then subjected to standard polymerase-ribozyme extension conditions. Nine time points were collected and analyzed by high-resolution PAGE. (A) For an A/U hairpin, triangles indicate the fraction extended of the PNK-repaired hairpin, crosses indicate the fraction extended of the PNK repaired hairpin in the presence of 10 mM PPi to determine the inhibition of the reaction by pyrophosphate, and open circles indicate the fraction extended of the 2′-3′-cyclic-phosphate-terminated hairpin that was repaired by the ribozyme in presence of 10 mM PPi. For hairpins terminated in G/C (B), U/A (C), or C/G (D), triangles indicate the fraction extended of the PNK-repaired hairpin, and open circles indicate the fraction extended of the 2′-3′-cyclic-phosphate-terminated hairpin that was repaired by addition of 10 mM PPi.

**Extended Data Fig. 10.**
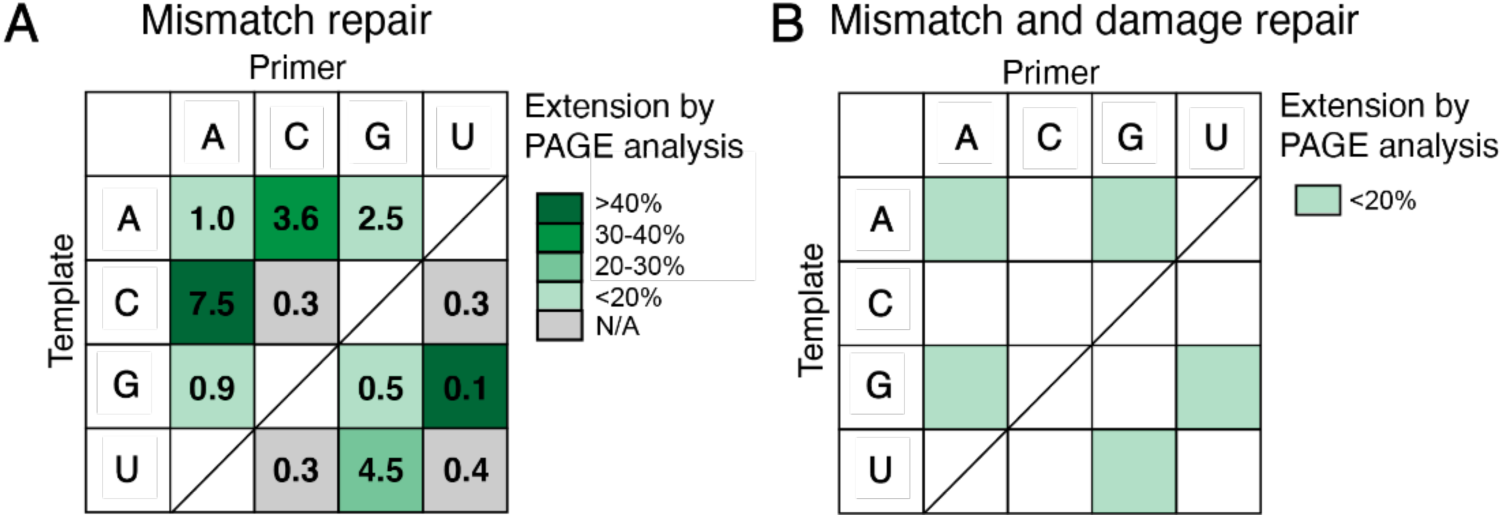
Trends in mismatch repair and damage and mismatch repair. (A) Grid showing the correlation between mismatch repair and overall mismatch extension. The box colors indicate what percentage of each mismatch is extended when repaired by the ribozyme in the presence of pyrophosphate, compared to a no-pyrophosphate control, as determined by PAGE analysis. The gray color indicates no significant difference observed in extension reactions in the presence/absence of pyrophosphate. The numbers correspond to the total fraction of mismatch repair derived from HTS data. (B) Grid showing mismatches in which both 2′-3′-cP phosphate damage and the mismatch could be repaired *via* ribozyme-catalyzed pyrophosphorolysis of the terminal nucleotide.

## Supplementary Information

### SI Note 1. Gel migration of cyclic-phosphate–terminated hairpins

Previous reports showed that treatment of 2′-3′-cP-terminated oligonucleotide sequences with PNK to remove the cyclic phosphate leads to a gel mobility retardation, and treatment of a 2′-3′-cyclic-phosphate-terminated sequence with acid to generate a linear monophosphate leads to a faster-migrating species.^52–54^ We consistently observed the opposite effect and have concluded that the anomalous migration is a property of the hairpin sequence, which differs structurally from the linear RNA sequences used in previously reported experiments. Following treatment with PNK, all 2′-3′-cP-terminated hairpin sequences exhibit faster gel migration. The relative gel mobility of the 2′-3′-cP-terminated hairpins treated with acid (which initially hydrolyzes just one of the phosphodiesters of the cyclic phosphate) shows a gel mobility retardation. This data confirmed that PNK treated hairpin sequences indeed migrate faster upon removal of the 2′-3′-cP (Extended Data Fig. 1). Many of the hairpin sequences when visualized by PAGE show two bands depending on the resolution of the experiment. Based on the gel migration of the acid treated 2′-3′-cP, we determined that these two bands arise from the difference between closed and open states of the terminal cyclic phosphate.

### SI Note 2. Analysis of HTS sequencing data for each relevant figure

Additional details on the HTS analysis for each figure:

All libraries were submitted for Illumina sequencing. See Supplementary Table 2 for explanation of datasets.

Illumina MiSeq Single Read 150 sequencing was used for tC19Z_MM_1, tC19Z_MM_2, tC19Z_MM_3, tC19Z_MM_4, tC19Z_MM_5, tC10Z_MM_6, tC19Z_MM_7, tC19Z_MM_8, tC19Z_MM_9, tC19Z_MM_10, tC19Z_MM_11, tC19Z_MM_12, tC19Z_MA_1, tC19Z_MA_2, tC19Z_MA_3, tC19Z_MA_4, tC19Z_MA_NoPPi_1, tC19Z_MA_NoPPi_2 Illumina MiSeq Paired End sequencing was used for tC19Z_LT_3_0, tC19Z_LT_3_2.5, tC19Z_LT_3_10, tC19Z_LT_4_0, tC19Z_LT_4_3, tC19Z_LT_4_10, tC19Z_LT_5_10, tC19Z_LT_5_2.5, tC19Z_LT_5_1, tC19Z_LT_5_0

A PhiX library was added into all samples and its sequences were first removed from all sequencing files using Bowtie2.

### Analysis for Figure 2d

CutAdapt was used to remove the adapters on first the 5′ and then the 3′-termini of the sequences. FastAptamer was used to count the sequences. FastAptamer search was then used to find all sequences that had the full hairpin present, including sequences in which the 3′-ends were either extended or shortened by up to 5 nucleotides. The total counts of these sequences were tabulated. FastAptamer search was then used to find sequences that corresponded to the starting hairpin, sequences that were −1, sequences that were extended by one nucleotide correctly, sequences that were extended by one nucleotide incorrectly, sequences that were extended by two nucleotides correctly, sequences that were extended by one nucleotide correctly and the second nucleotide incorrectly, etc., up to five nucleotides. These sequences were then visually inspected and manually curated. The counts of the remaining sequences were tabulated. The extension percentages shown in Figure 2d were calculated by generating percentages of the total sequences for each extension length regardless of extended nucleotide identity.

### Analysis for Figure 2e

CutAdapt was used to remove the adapters on first the 5′ and then the 3′-termini of the sequences. FastAptamer was used to count the sequences. FastAptamer search was then used to find all sequences that had the hairpin present, including sequences in which the 3′-ends were either extended or shortened by up to 5 nucleotides. The total counts of these sequences were tabulated. FastAptamer search was then used to find sequences that corresponded to the starting hairpin, sequences that were −1, sequences that were extended by one nucleotide correctly, sequences that were extended by one nucleotide incorrectly, sequences that were extended by two nucleotides correctly, sequences that were extended by one nucleotide correctly and the second nucleotide incorrectly, etc., up to five nucleotides. These sequences were then visually inspected and manually curated. The counts of the remaining sequences were tabulated. The extension percentages shown in Figure 2e were calculated by generating percentages of just sequences, in which the nucleotides were extended correctly as shown in the figure. The same normalization factor as used in Figure 2e was applied.

### Figure 2f

CutAdapt was used to remove the adapters on first the 5′ and then the 3′-termini of the sequences. FastAptamer was used to count the sequences. FastAptamer search was then used to find all sequences that had the hairpin present, including sequences in which the 3′-ends were either extended or shortened by up to 5 nucleotides. The total counts of these sequences were tabulated. FastAptamer search was then used to find sequences that corresponded to the starting hairpin, sequences that were −1, sequences that were extended by one nucleotide correctly, sequences that were extended by one nucleotide incorrectly, sequences that were extended by two nucleotides correctly, sequences that were extended by one nucleotide correctly and the second nucleotide incorrectly, etc., up to five nucleotides. These sequences were then visually inspected and manually curated. The counts of the remaining sequences were tabulated. The percentages shown in Figure 2f were calculated by tabulating all sequences that had the mismatched nucleotide repaired and extended from zero to five nucleotides for each mismatched sequence.

### Figure 2g

CutAdapt was used to remove the adapters on first the 5′ and then the 3′-termini of the sequences. FastAptamer was used to count the sequences. FastAptamer search was then used to find all sequences that had the hairpin present, including sequences in which the 3′-ends were either extended or shortened by up to 5 nucleotides. The total counts of these sequences were tabulated. FastAptamer search was then used to find sequences that corresponded to the starting hairpin, sequences that were −1, sequences that were extended by one nucleotide correctly, sequences that were extended by one nucleotide incorrectly, sequences that were extended by two nucleotides correctly, sequences that were extended by one nucleotide correctly and the second nucleotide incorrectly, etc., up to five nucleotides. These sequences were then visually inspected and manually curated. The counts of the remaining sequences were tabulated. The percentages shown in the black bars in Figure 2g were calculated by tabulating the sequences that were not repaired and the percentage that were subsequently extended correctly was calculated. The grey bars in Figure 2g were calculated by tabulating the sequences that were repaired and the percentage that were subsequently extended correctly was calculated.

### Figure 3b and 3c

CutAdapt was used to remove the adapters and primer regions on first the 5′ and then the 3′-end of the sequences. Sequences were counted in FastAptamer using fastaptamer_count. Sequences containing the primer were extracted from data for analysis in FastAptamer using fastaptamer_search for each sequence and subsequently visually aligned and curated using Jalview. Sequences that were not extended, ligation artifacts, and sequencing artifacts were removed. Jalview was used to view the aligned sequences. The sequence lengths were verified visually. This procedure was repeated for each condition shown in the figure.

### Figure 3d and 3e

CutAdapt was used to remove the adapters and primer regions on first the 5′ and then the 3′-end of the sequences. Sequences were counted in FastAptamer using fastaptamer_count. Sequences containing the primer were extracted from data for analysis in FastAptamer using fastaptamer_search for each sequence and subsequently visually aligned and curated using Jalview. Sequences that were not extended, ligation artifacts, and sequencing artifacts were removed. The fidelity was determined by visually inspecting the sequences in Jalview and tabulating the number of incorrectly incorporated nucleotides. The percentage fidelity was determined as the average of the fidelity at each position.

### Figure 4c-g

CutAdapt was used to remove the adapters on first the 5′ and then the 3′-termini of the sequences. FastAptamer was used to count the sequences. FastAptamer search was then used to find all sequences that had the hairpin present, including sequences in which the 3′-ends were either extended or shortened by up to 5 nucleotides. The total counts of these sequences were tabulated. FastAptamer search was then used to find sequences that corresponded to the starting hairpin, sequences that were −1, sequences that were extended by one nucleotide correctly, sequences that were extended by one nucleotide incorrectly, sequences that were extended by two nucleotides correctly, sequences that were extended by one nucleotide correctly and the second nucleotide incorrectly, etc., up to five nucleotides. These sequences were then visually inspected and manually curated. The counts of the remaining sequences were tabulated. The extension percentages shown in Figure 4c-g were calculated by generating percentages of the total sequences for each extension length.

### Extended Data Fig. 4

CutAdapt was used to remove the adapters on first the 5′ and then the 3′-end of the sequences. FastAptamer was used to count the sequences. FastAptamer search was then used to find all sequences that had the hairpin present including sequences where the 3′-end was either extended or shortened up to 5 nucleotides. The total counts of these sequences were tabulated. FastAptamer search was then used to find sequences that corresponded to sequences that were −1, −2, −3, and −4. These sequences were then visually inspected and manually curated. The counts of the remaining sequences were tabulated. The percentages shown in Extended Data Fig. 4 were calculated by generating percentages of sequences that had 1-4 nucleotides removed.

### Extended Data Fig. 5

CutAdapt was used to remove the adapters on first the 5′ and then the 3′-termini of the sequences. FastAptamer was used to count the sequences. FastAptamer search was then used to find all sequences that had the hairpin present, including sequences in which the 3′-ends were either extended or shortened by up to 5 nucleotides. The total counts of these sequences were tabulated. FastAptamer search was then used to find sequences that corresponded to the starting hairpin, sequences that were −1, sequences that were extended by one nucleotide correctly, sequences that were extended by one nucleotide incorrectly, sequences that were extended by two nucleotides correctly, sequences that were extended by one nucleotide correctly and the second nucleotide incorrectly, etc., up to five nucleotides. These sequences were then visually inspected and manually curated. The counts of the remaining sequences were tabulated. The extension percentages shown in the left graph of each panel in Extended Data Fig. 5 were calculated by generating percentages of the total sequences for each extension length regardless of extended nucleotide identity. The percentages were then normalized to add to 100%. The right graph of each panel in Extended Data Fig. 5 shows the percentage of the sequences in the left panel at each length where the mismatch was corrected.

### Extended Data Fig. 6

CutAdapt was used to remove the adapters on first the 5′ and then the 3′-termini of the sequences. FastAptamer was used to count the sequences. FastAptamer search was then used to find all sequences that had the hairpin present, including sequences in which the 3′-ends were either extended or shortened by up to 5 nucleotides. The total counts of these sequences were tabulated. FastAptamer search was then used to find sequences that corresponded to the starting hairpin, sequences that were −1, sequences that were extended by one nucleotide correctly, sequences that were extended by one nucleotide incorrectly, sequences that were extended by two nucleotides correctly, sequences that were extended by one nucleotide correctly and the second nucleotide incorrectly, etc., up to five nucleotides. These sequences were then visually inspected and manually curated. The counts of the remaining sequences were tabulated. Extension percentages shown in Extended Data Fig. 6 were calculated for each sequence shown as a fraction of the total hairpin containing sequences.

### Supplementary Fig. 2

CutAdapt was used to remove the adapters and primer regions on first the 5′ and then the 3′-end of the sequences, including the seven random nucleotides. Sequences were counted in FastAptamer using fastaptamer_count. Sequences containing the primer were extracted from data for analysis in FastAptamer using fastaptamer_search for each sequence and subsequently visually aligned and curated using Jalview. Sequences that were not extended, ligation artifacts, and sequencing artifacts were removed. The sequences were visually inspected, manually aligned, and sequences that did not align to the start of the sequence were removed. The sequences were exported to Adobe Illustrator, the reference sequence was colored green and mismatches were colored blue.

**Supplementary Fig. 1.**
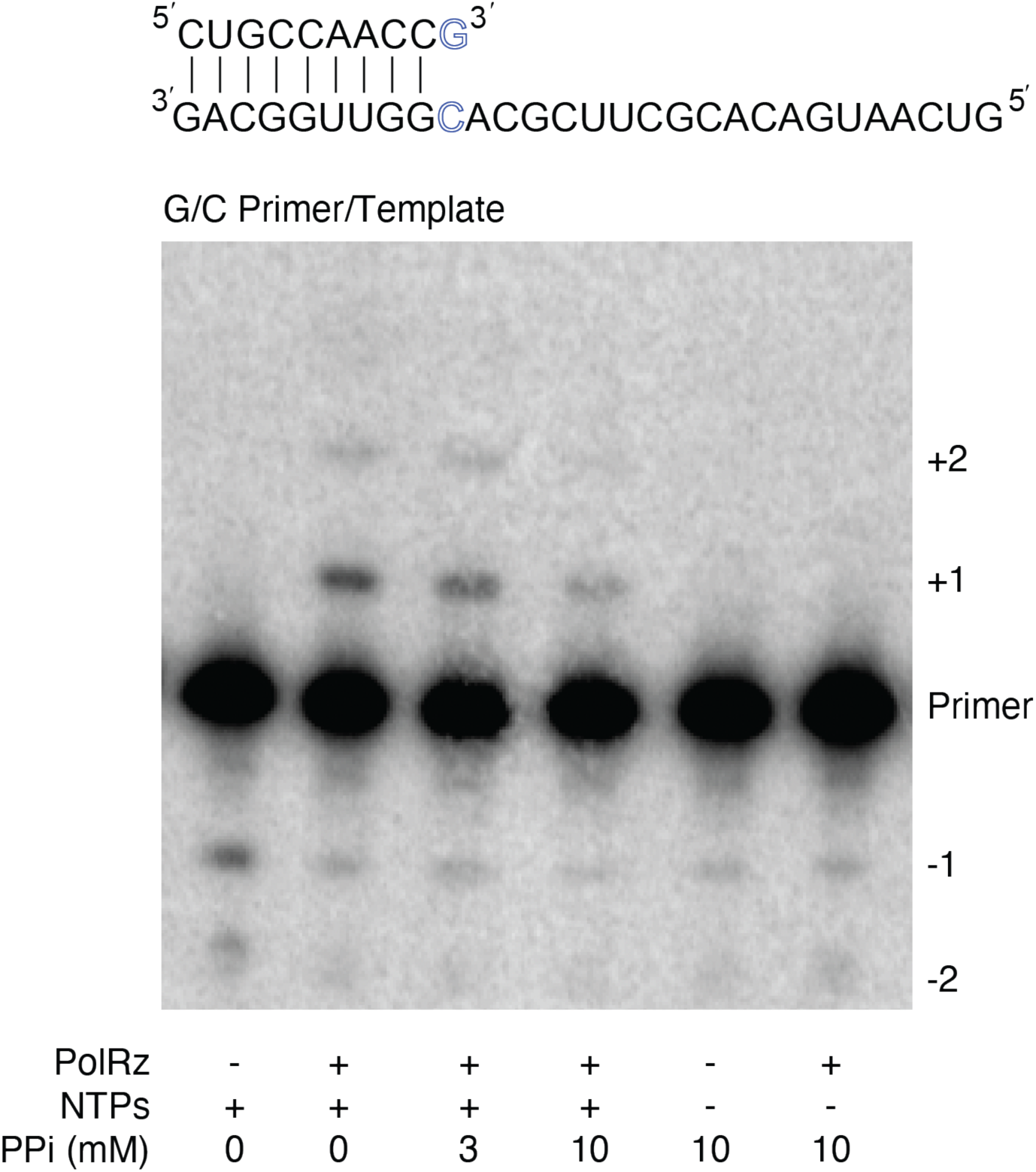
Pyrophosphate inhibits ribozyme mediated extension. A primer/template sequence is extended by the ribozyme under standard conditions. The extension is inhibited when pyrophosphate is also added to the reaction (see also Extended Data Fig. 3). However, addition of pyrophosphate either with or without the tC19Z polymerase ribozyme does not result in any additional bands corresponding to RNA degradation or significant observable pyrophosphorolysis.

**Supplementary Fig. 2.**
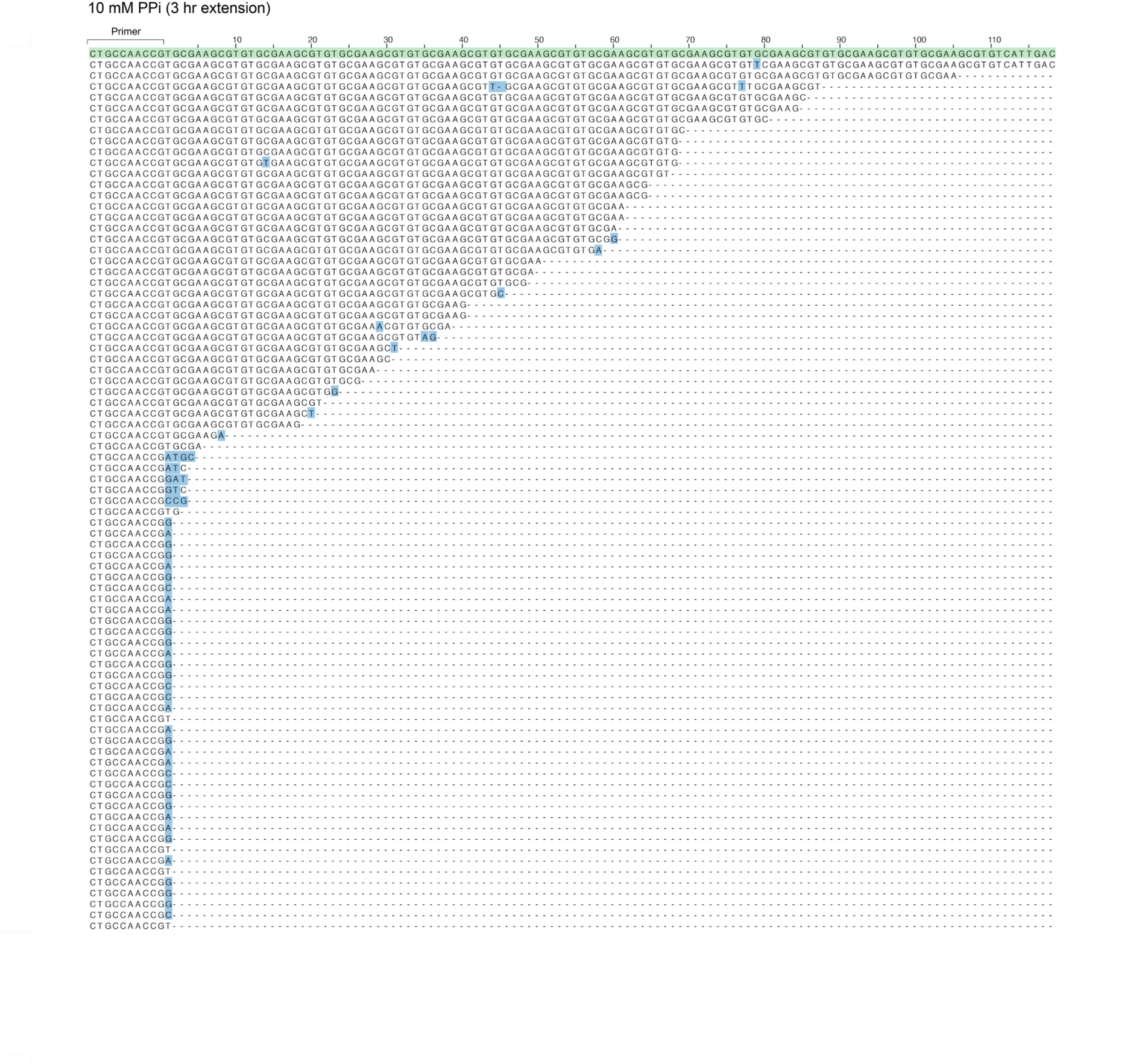
Alignment of long extension sequences from HTS. Sequences corresponding to extensions present in the HTS data generated from the three-hour polymerization reaction and subsequent sequencing described in Figure 3B. Many of the sequences that are not full length terminate in one or more mismatches. The primer region is shown in the bracket, and the expected full-length sequence the HTS data was aligned to is highlighted in green. Mismatches are highlighted in blue.

**Supplementary Fig. 3.**
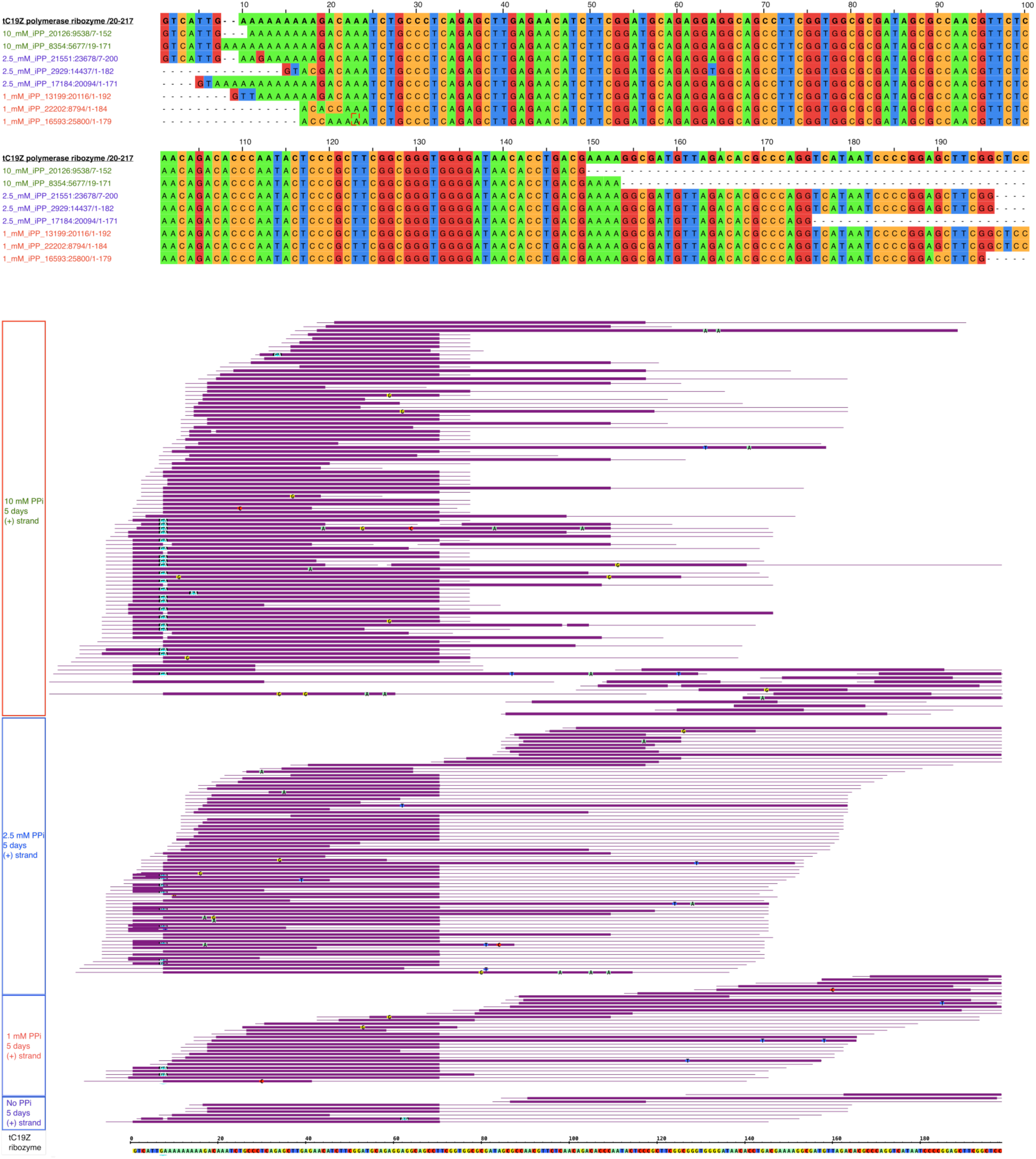
tC19Z sequences present in HTS analysis of long extension reactions. (Top) Sequences of cDNAs generated by 3’-terminal ligation of the RNA strand and, after reverse transcription, 3’-terminal ligation of the cDNA strand. The high-throughput sequencing forward adapter was ligated to the 3’-terminus of the RNA. The cDNAs shown matching the tC19Z ribozyme (top sequence) therefore report generation of RNAs antisense to the ribozyme during the extension reactions (data shown are for analysis of 5-day experiments). RNAs corresponding to long segments of tC19Z were observed only in experiments containing pyrophosphate. The pyrophosphate concentrations are annotated on the left. The longest copies of the ribozyme lack just 4 nucleotides (2%) from the ribozyme termini. (Bottom) All cDNA sequences matching the sense strand of the tC19Z ribozyme generated under 0-, 1-, 2.5-, and 10-mM pyrophosphate conditions. No long sequences were observed in samples lacking pyrophosphate. Thick lines represent the ribozyme matching segments, whereas the thin lines represent segments of other sequences, such as ligation and sequencing adapters.

**Supplementary Fig. 4.**
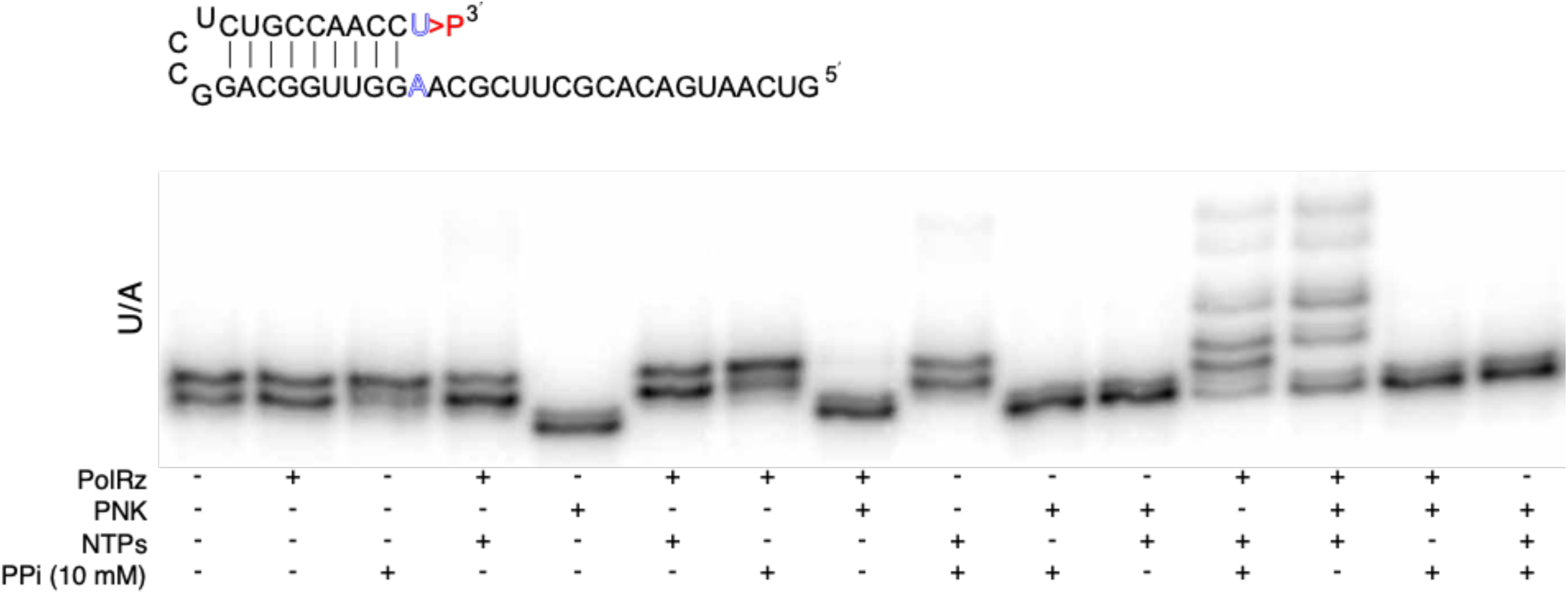
Damage repair reactions with a full set of controls for the U/A-terminated hairpin substrate. A U/A hairpin was allowed to react under standard extension conditions with variations shown by + and − symbols below the gel image.

**Supplementary Fig. 5.**
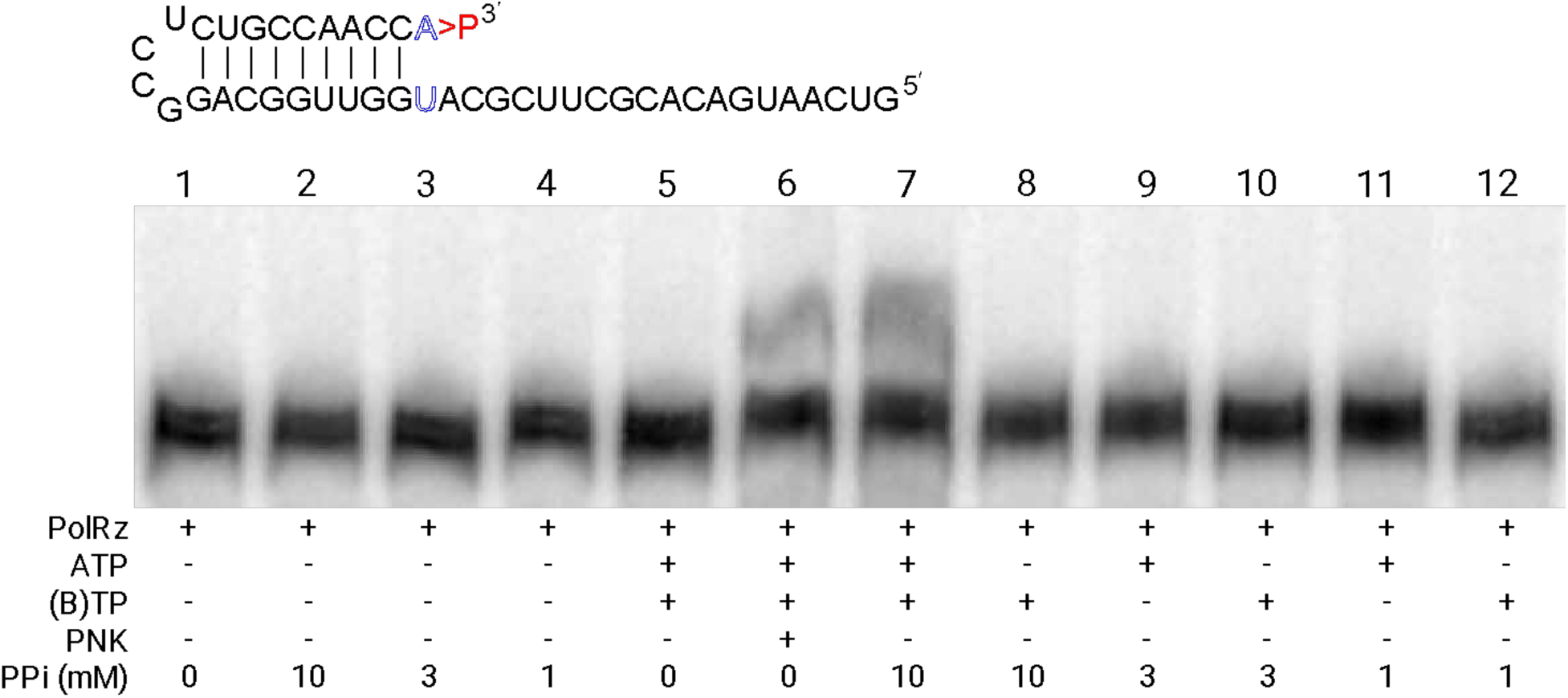
Omission of a single nucleotide prevents extension of a repaired A/U hairpin. A ^32^P-labeled A/U hairpin was extended under standard reaction conditions with variations indicated by the + and – symbols below each reaction resolved by PAGE. The left 5 lanes are negative controls, that lack any NTP; lane 6 is a positive control (PNK-repaired hairpin extension); and lane 7 is the pyrophosphate-facilitated repair reaction at 10 mM pyrophosphate. Lane 8 indicates that upon addition of all nucleotides except ATP (indicated by (B)TP (all, except ATP)), no extension is observed, strongly suggesting that the pyrophosphate-facilitated repair removes adenosine 5′ monophosphate-2′-3′-cyclic phosphate from the damaged terminus and requires ATP to incorporate the first nucleotide. Lanes 9–12 show that the repair does not occur at subsaturating concentrations of pyrophosphate.

**Supplementary Fig. 6.**
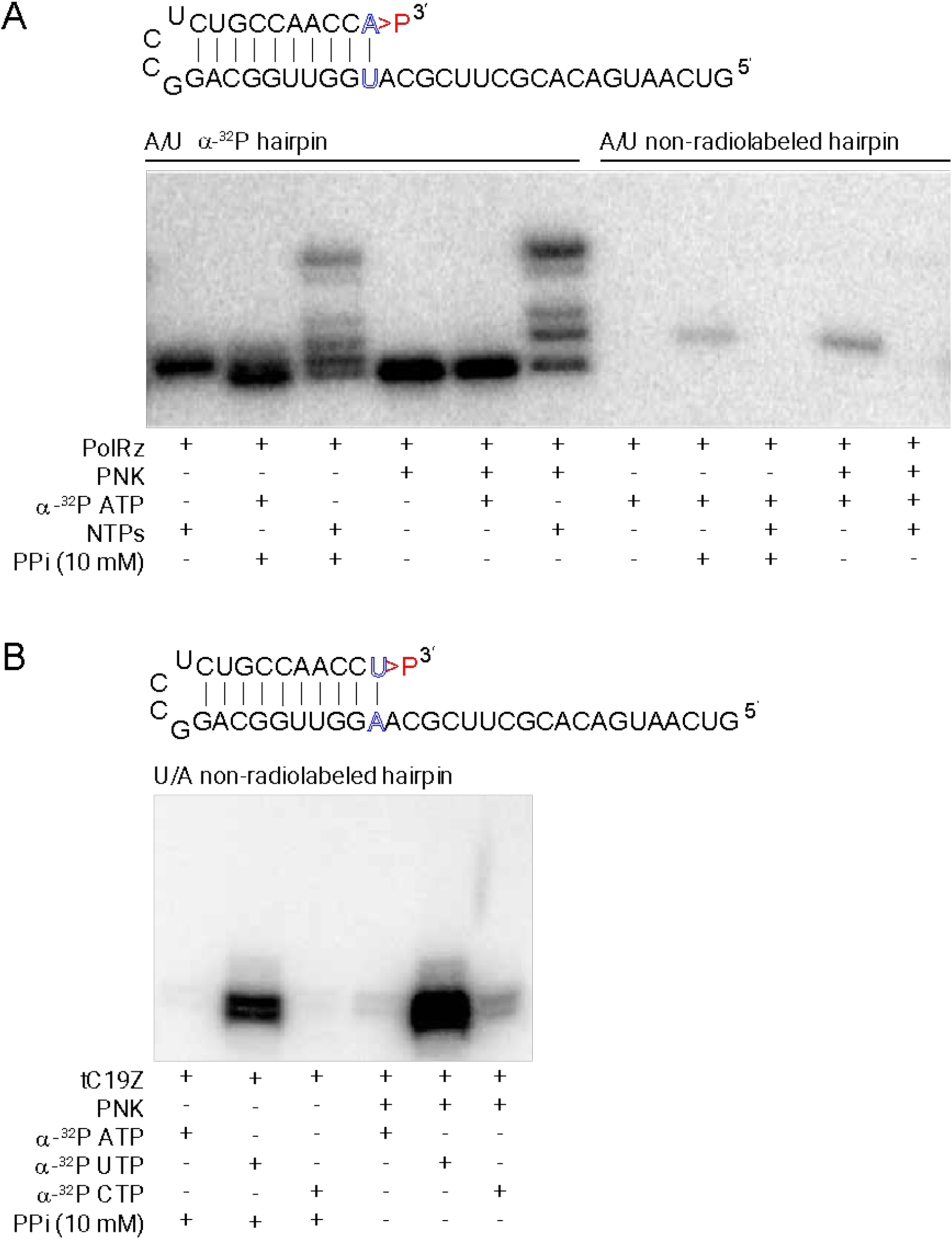
Incorporation of α-^32^P-labeled nucleotides into repaired, non-labeled hairpins. (A) An A/U hairpin was prepared with or without internal ^32^P, as indicated above the PAGE gel image. The hairpin was then subjected to repair and extension under standard reaction conditions, except that *α*-^32^P-ATP was added to some reactions. In the case of the non-labeled hairpin, the ^32^P-AMP is only incorporated if the sequence has been repaired either by pyrophosphate-facilitated reaction or by PNK. (B) A U/A hairpin was transcribed without *α*-^32^P-ATP and was subjected to standard extension conditions but using *α*-^32^P-rNTP (indicated) in place of the standard NTPs. Different *α*-^32^P-rNTPs are utilized by the tC19Z polymerase ribozyme with different efficiencies, as shown by the varying intensities of the bands. ^32^P-UMP is incorporated most efficiently because the first nucleotide on the template sequence is an adenosine. However, misincorporation is seen for the other nucleotides, and a second nucleotide appears to be added in presence of UTP. No ^32^P labeling is observed in control reactions that lack PNK or pyrophosphate (as shown in panel A).

**Supplementary Fig. 7.**
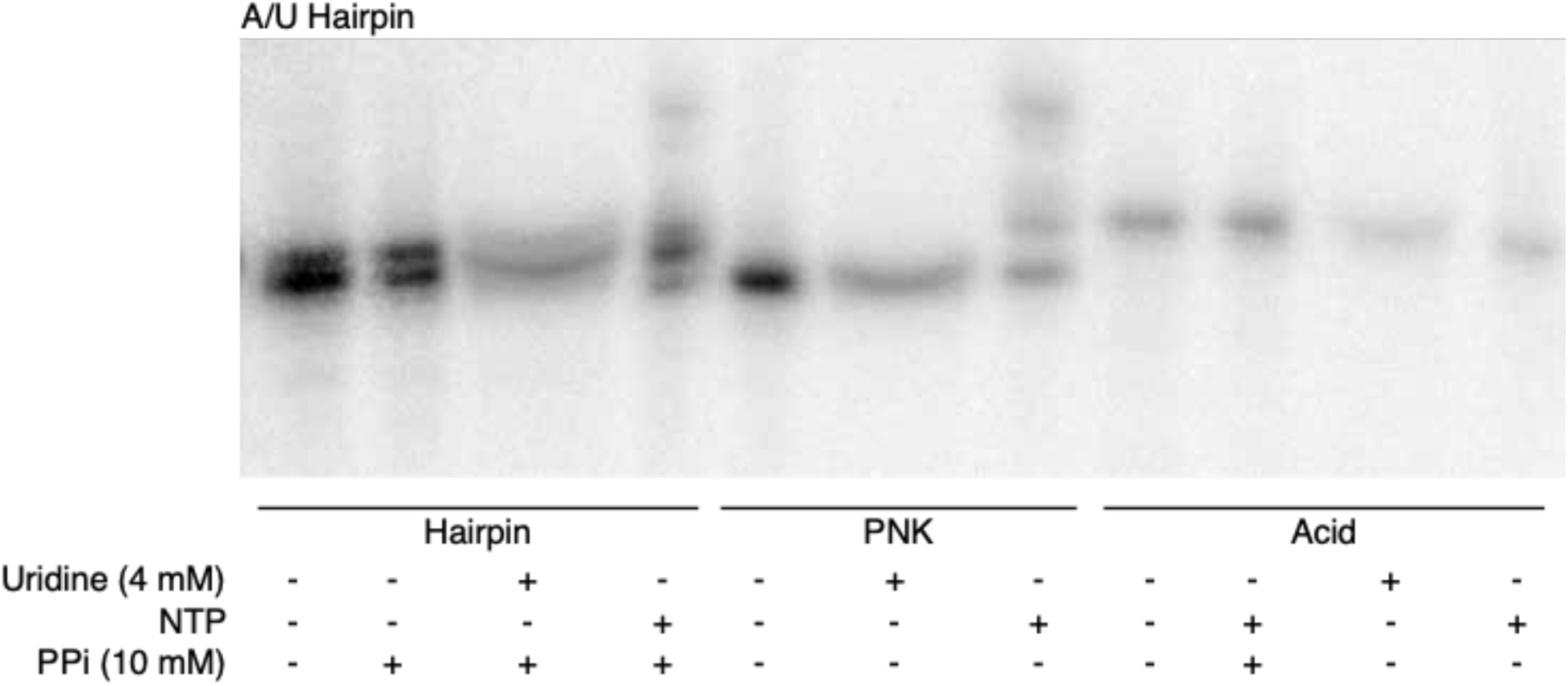
The mechanism of repair does not involve cyclic-phosphate opening by pyrophosphate. Pyrophosphate and uridine were added to a damaged A/U hairpin to determine whether the cyclic phosphate could be opened by pyrophosphate to form a triphosphate and allow for subsequent addition of uridine (lane 3). No additional bands were observed on the PAGE gel, indicating that this is not the mechanism of repair. Uridine was added to PNK-repaired or acid-treated hairpins as positive and negative controls, respectively.

**Supplementary Fig. 8.**
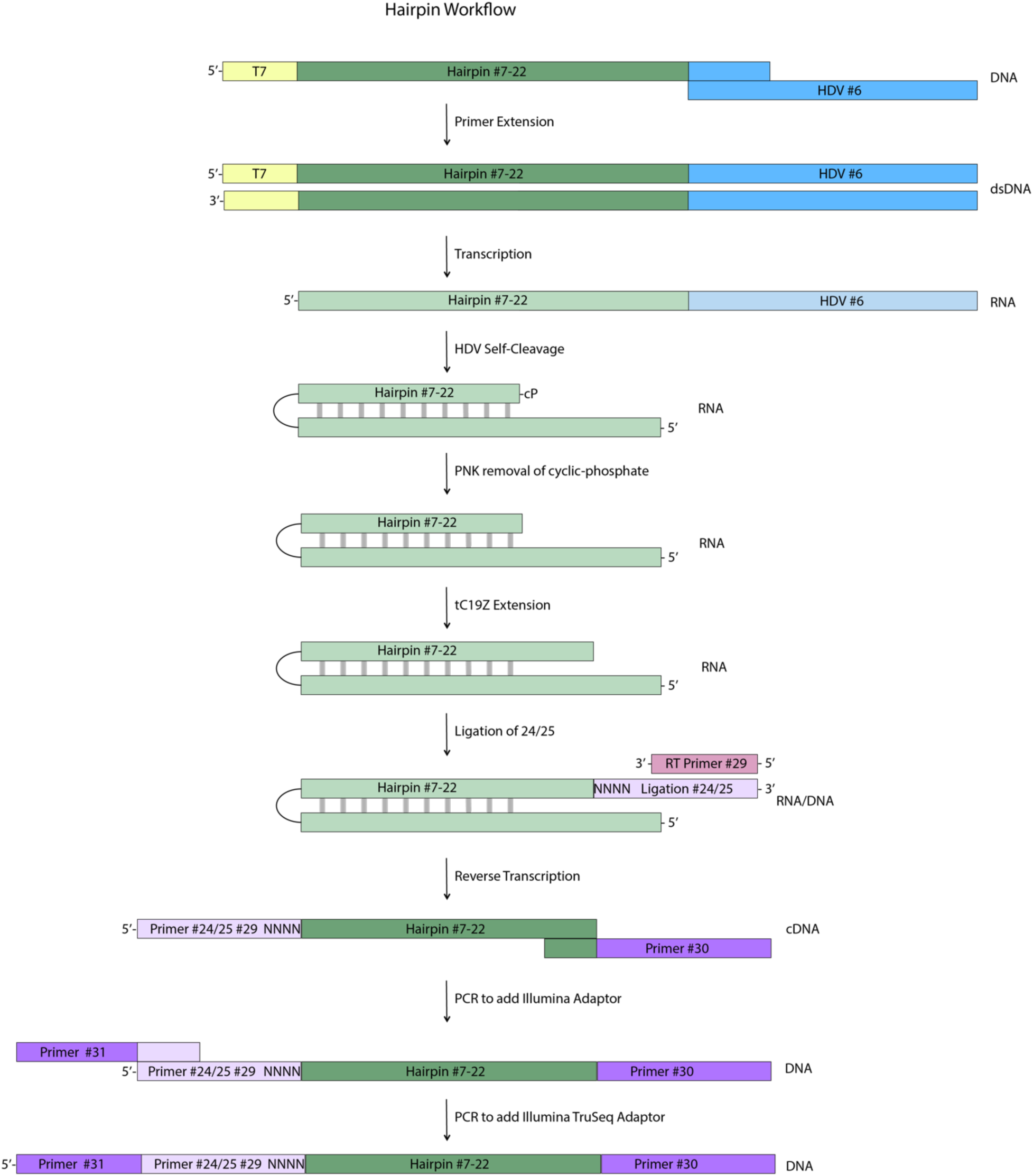
Workflow for HTS analysis of hairpin substrates of the polymerase ribozyme. The hairpins were first prepared by primer extension of an drz-Mbac-1 HDV-like ribozyme sequence (#6) and a hairpin sequence (#7–#22). This DNA construct with the ribozyme on the 3′-end of the hairpin was transcribed into RNA where the HDV-like ribozyme co-transcriptionally cleaves, leaving a hairpin with a 2′-3′-cyclic phosphate. For mismatch experiments, that cyclic phosphate can be removed during incubation with PNK. The hairpins are then extended by the tC19Z polymerase ribozyme. Following this reaction, the RNA is ligated to a DNA oligo with a randomized 5′-end (#24/#25) and the resulting construct is reverse-transcribed with RT primer #29. The Illumina adapters (#30 and #31) are then added via PCR prior to sequencing.

**Supplementary Table 1.**
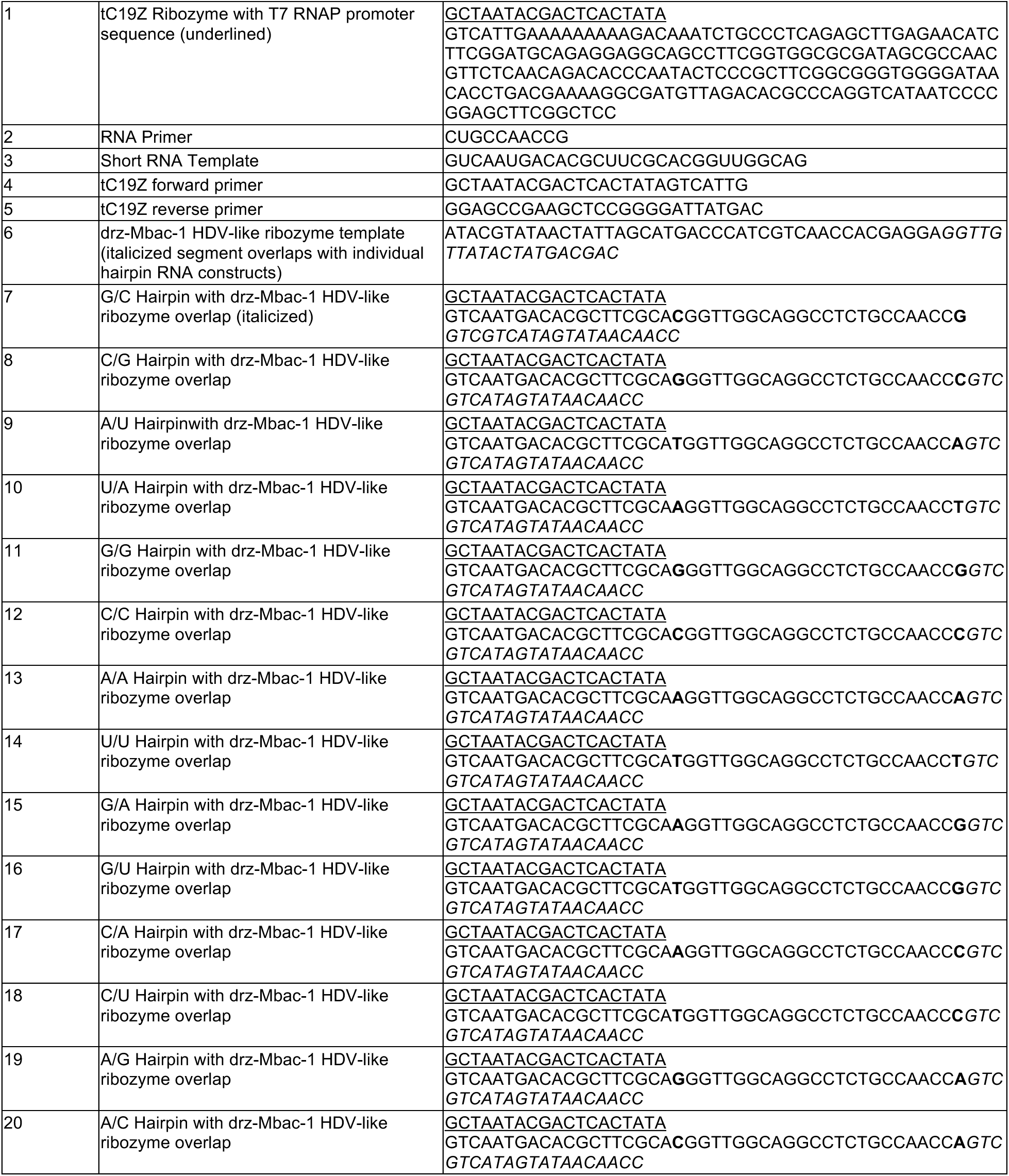
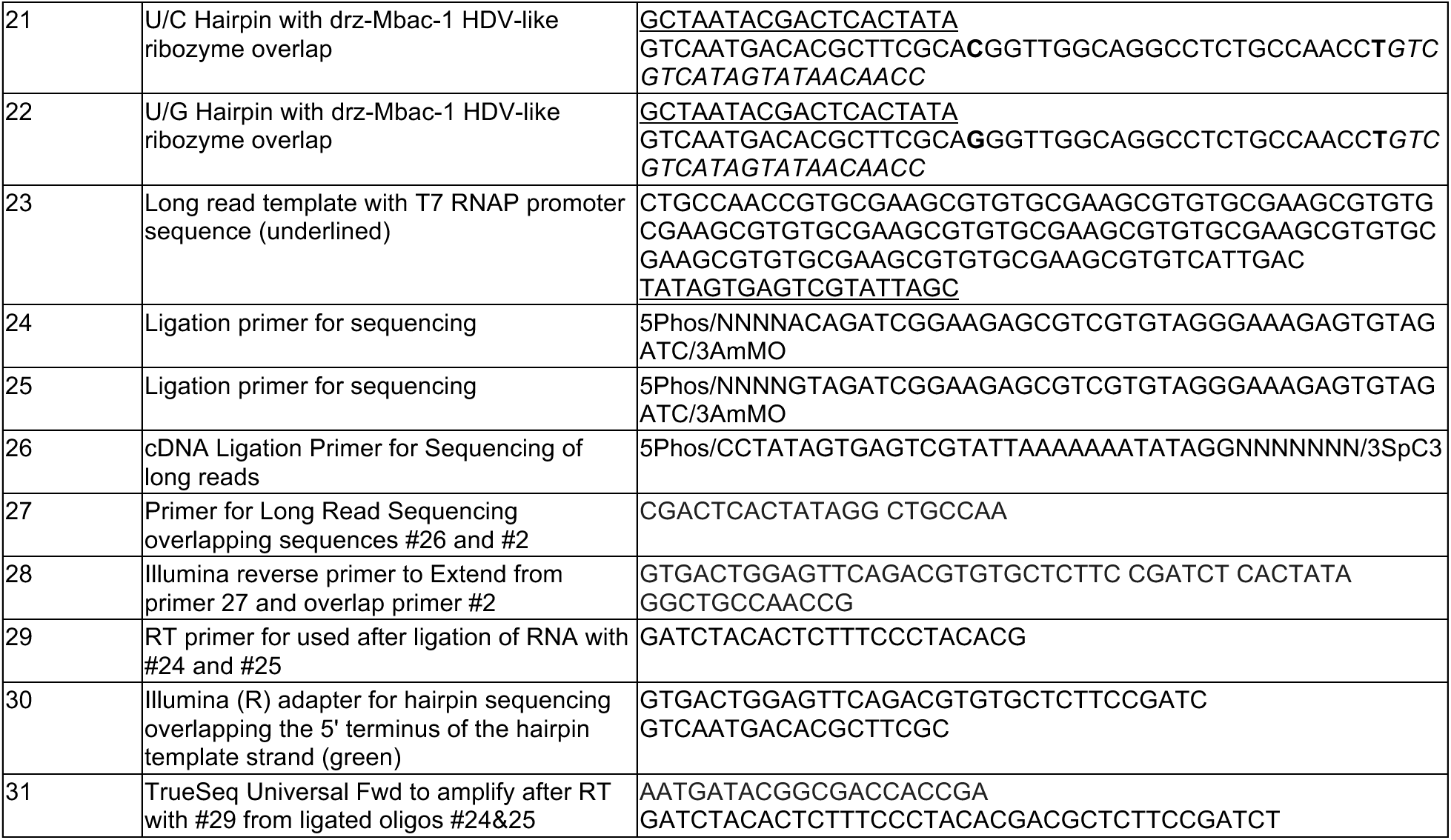
DNA and RNA sequences. T7 RNA polymerase promoter sequences are underlined. HDV overlap sequences in the hairpin constructs are italicized. The terminal match or mismatch in the hairpin sequences is bolded.

**Supplementary Table 2.** Correlation of figures to deposited sequencing data.

| Dataset | Main Figures | Extended Data and SI Figures |
| --- | --- | --- |
| tC19Z_MM_1 | 2f (U/G), 2g (U/G) | Extended Data Fig. 4C (U/G),<br>Extended Data Fig. 5F |
| tC19Z_MM_2 | 2f (G/U), 2g (G/U) | Extended Data Fig. 4C (G/U) |
| tC19Z_MM_3 | 2d, 2e, 2f (A/C), 2g (A/C) | Extended Data Fig. 4C (A/C),<br>Extended Data Fig. 5D |
| tC19Z_MM_4 | 2f (C/A), 2g (C/A) | Extended Data Fig. 4C (C/A),<br>Extended Data Fig. 5E |
| tC19Z_MM_5 | 2f (A/A), 2g (A/A) | Extended Data Fig. 4C (A/A) |
| tC19Z_MM_6 | 2f (A/G), 2g (A/A) | Extended Data Fig. 4C (A/G),<br>Extended Data Fig. 5B |
| tC19Z_MM_7 | 2f (G/G), 2g (G/G) | Extended Data Fig. 4C (G/G),<br>Extended Data Fig. 5A |
| tC19Z_MM_8 | 2f (G/A), 2g (G/A) | Extended Data Fig. 4C (G/G),<br>Extended Data Fig. 5C |
| tC19Z_MM_9 | 2f (U/U), 2g (U/U) | Extended Data Fig. 4C (U/U) |
| tC19Z_MM_10 | 2f (U/C), 2g (U/C) | Extended Data Fig. 4C (U/C),<br>Extended Data Fig. 6 |
| tC19Z_MM_11 | 2f (C/C), 2g (C/C) | Extended Data Fig. 4C (C/C),<br>Extended Data Fig. 5H |
| tC19Z_MM_12 | 2f (C/U), 2g (C/U) | Extended Data Fig. 4C (C/U),<br>Extended Data Fig. 5G |
| tC19Z_MA_1 | 4e | Extended Data Fig. 4B (U/A) |
| tC19Z_MA_2 | 4c, 4d |  |
| tC19Z_MA_3 | 4f |  |
| tC19Z_MA_4 | 4g | Extended Data Fig. 4B (C/G) |
| tC19Z_MA_NoPPi_1 |  | Extended Data Fig. 4A (U/A) |
| tC19Z_MA_NoPPi_2 |  | Extended Data Fig. 4A (C/G) |
| tC19Z_LT_3_0 | 3b, 3c (0 mM PPi) | Supplementary Fig. 2 |
| tC19Z_LT_3_2.5 | 3b, 3c (2.5 mM PPi) |  |
| tC19Z_LT_3_10 | 3b, 3c (10 mM PPi) | Supplementary Fig. 2 |
| tC19Z_LT_4_0 | 3b, 3d (0 mM PPi) |  |
| tC19Z_LT_4_3 | 3b, 3d (3 mM PPi) |  |
| tC19Z_LT_4_10 | 3b, 3d (10 mM PPi) |  |
| tC19Z_LT_0_0 | 3b (0 mM PPi) |  |

These datasets are available in NLM under SRA PRJNA1174777.

**Supplementary Table 3:** Sequencing data in Figure 3.

| Dataset | Total seq. | % Seq. with primer | % Extended |
| --- | --- | --- | --- |
| 0 hours, 0 mM PPI | 50825 | 14.7 | 0.08 |
| 3 hours, 0 mM PPI | 101862 | 30.9 | 0.83 |
| 3 hours, 2.5 mM PPI | 56588 | 37.9 | 0.77 |
| 3 hours, 10 mM PPI | 42930 | 35.6 | 0.52 |
| 4 days, 0 mM PPI | 140423 | 25.2 | 1.11 |
| 4 days, 3 mM PPI | 65769 | 17.1 | 1.39 |
| 4 days, 10 mM PPI | 26457 | 22.3 | 1.13 |

## Supplementary Information: Full Length Gels

Boxes have been drawn around relevant lanes.

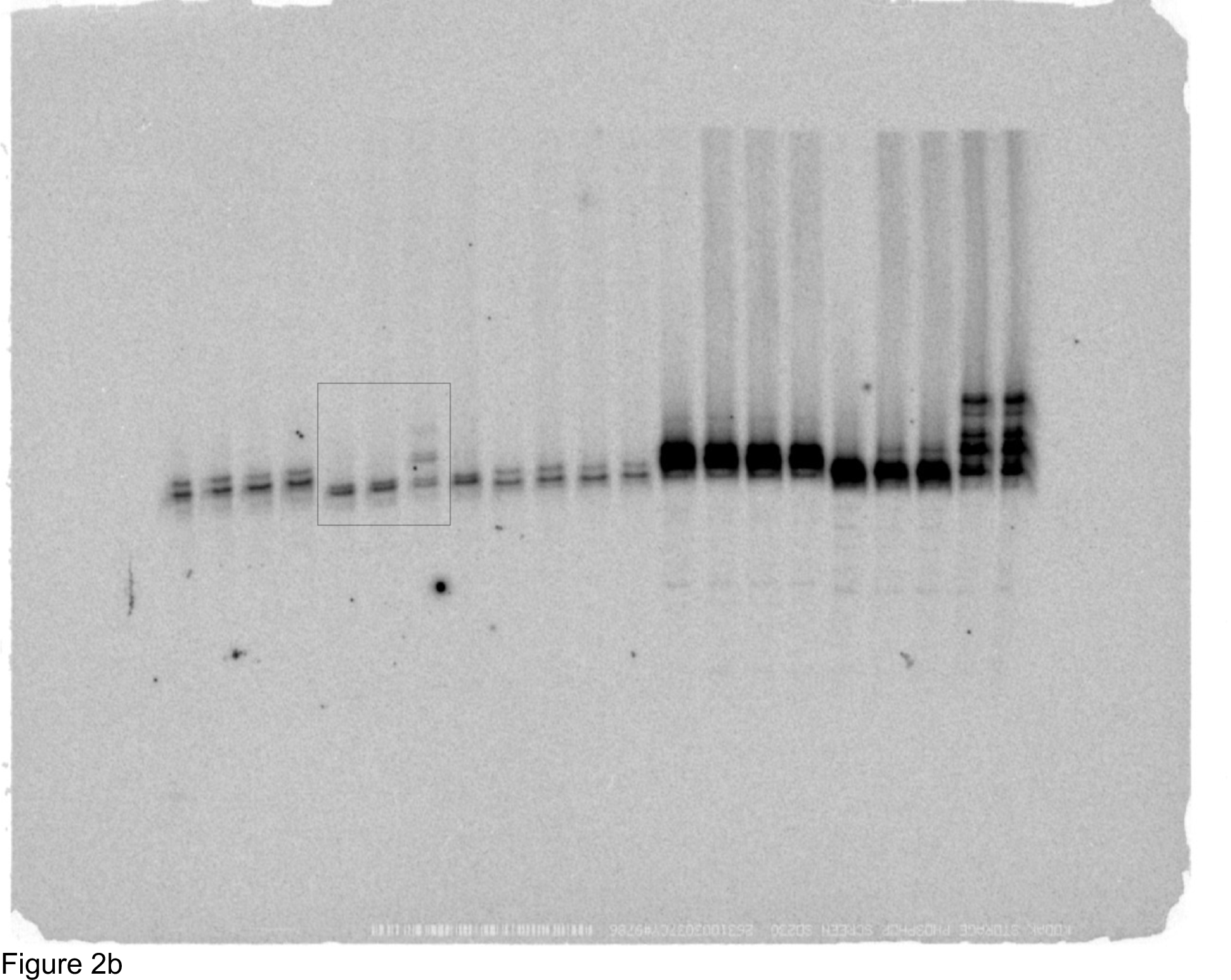

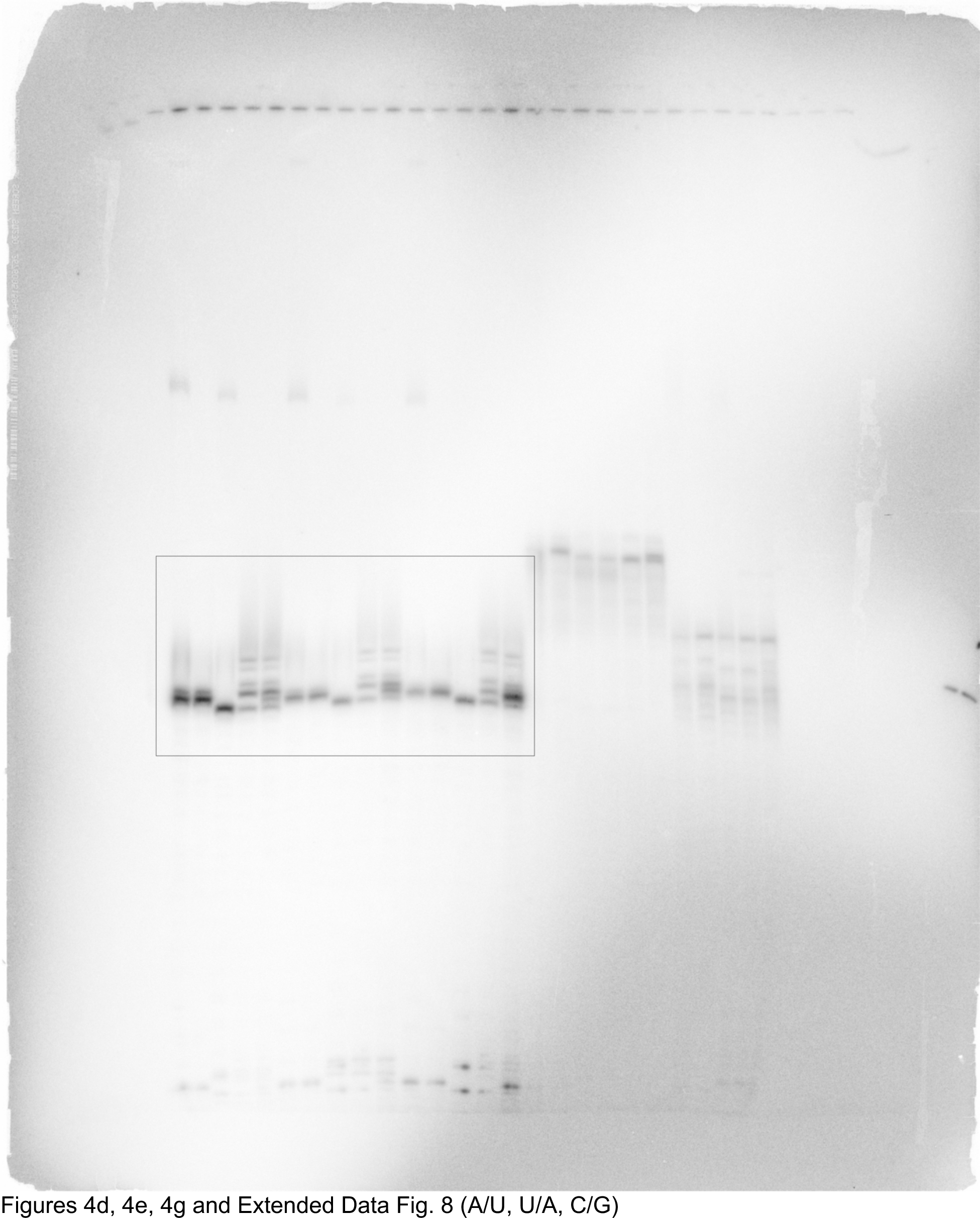

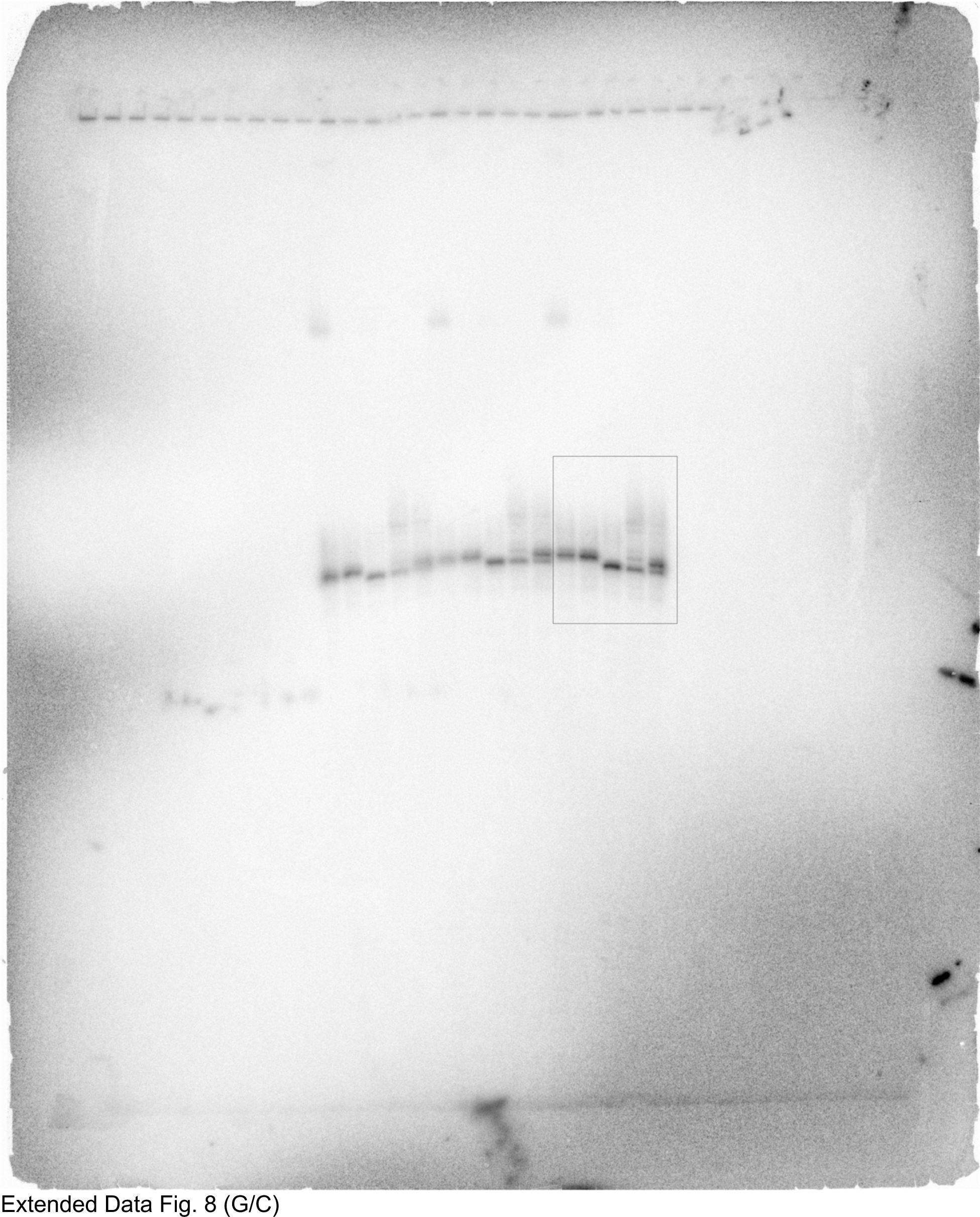

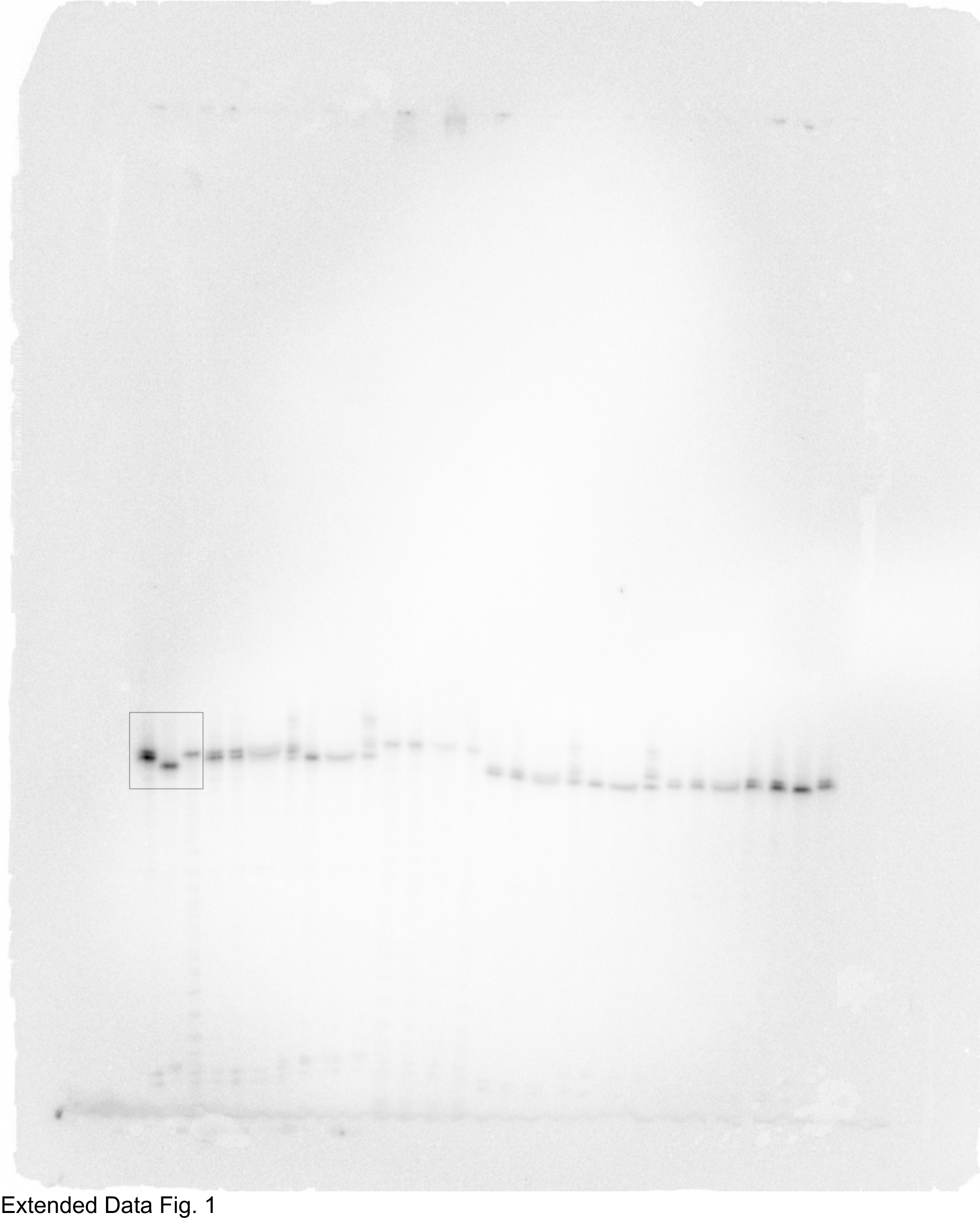

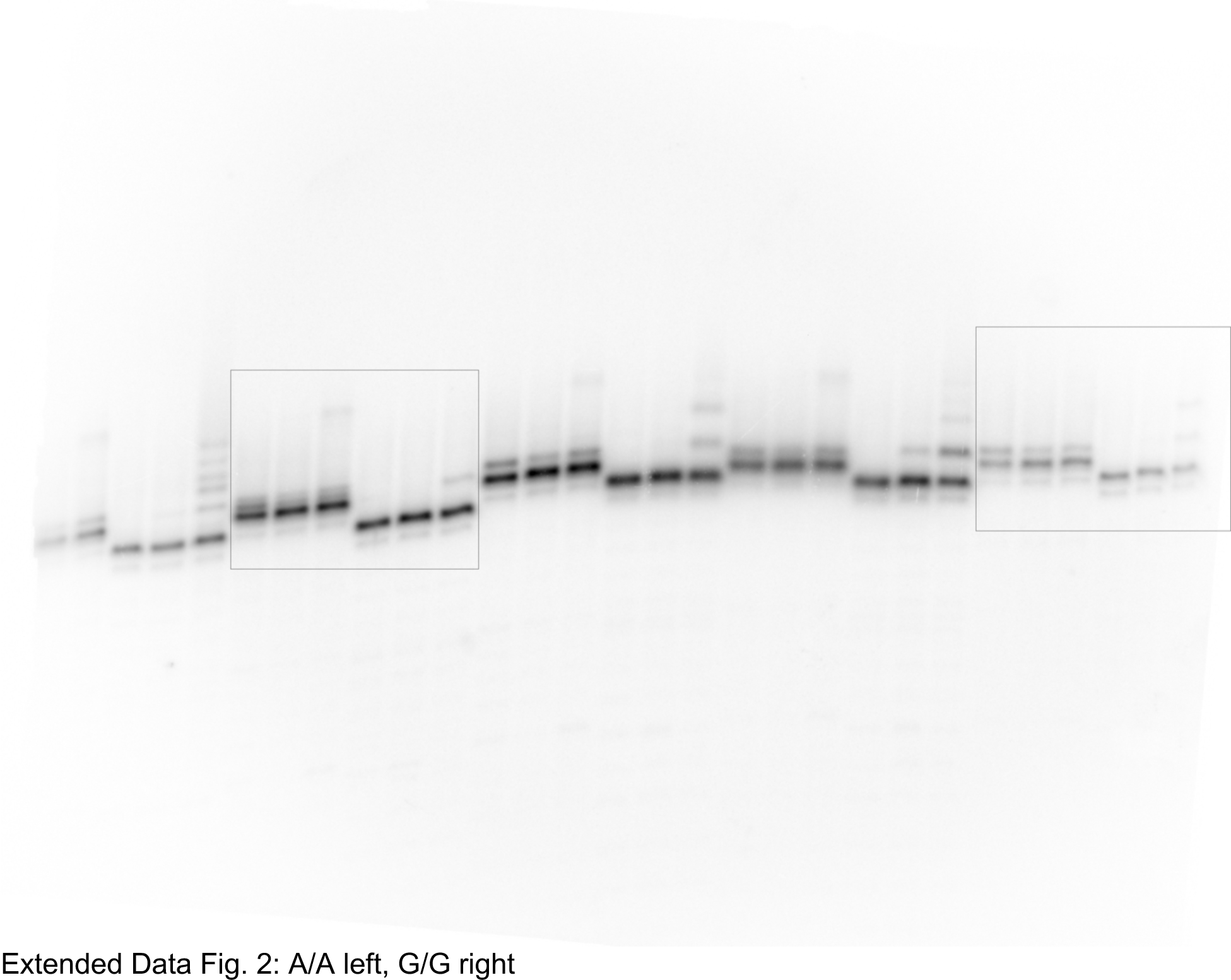

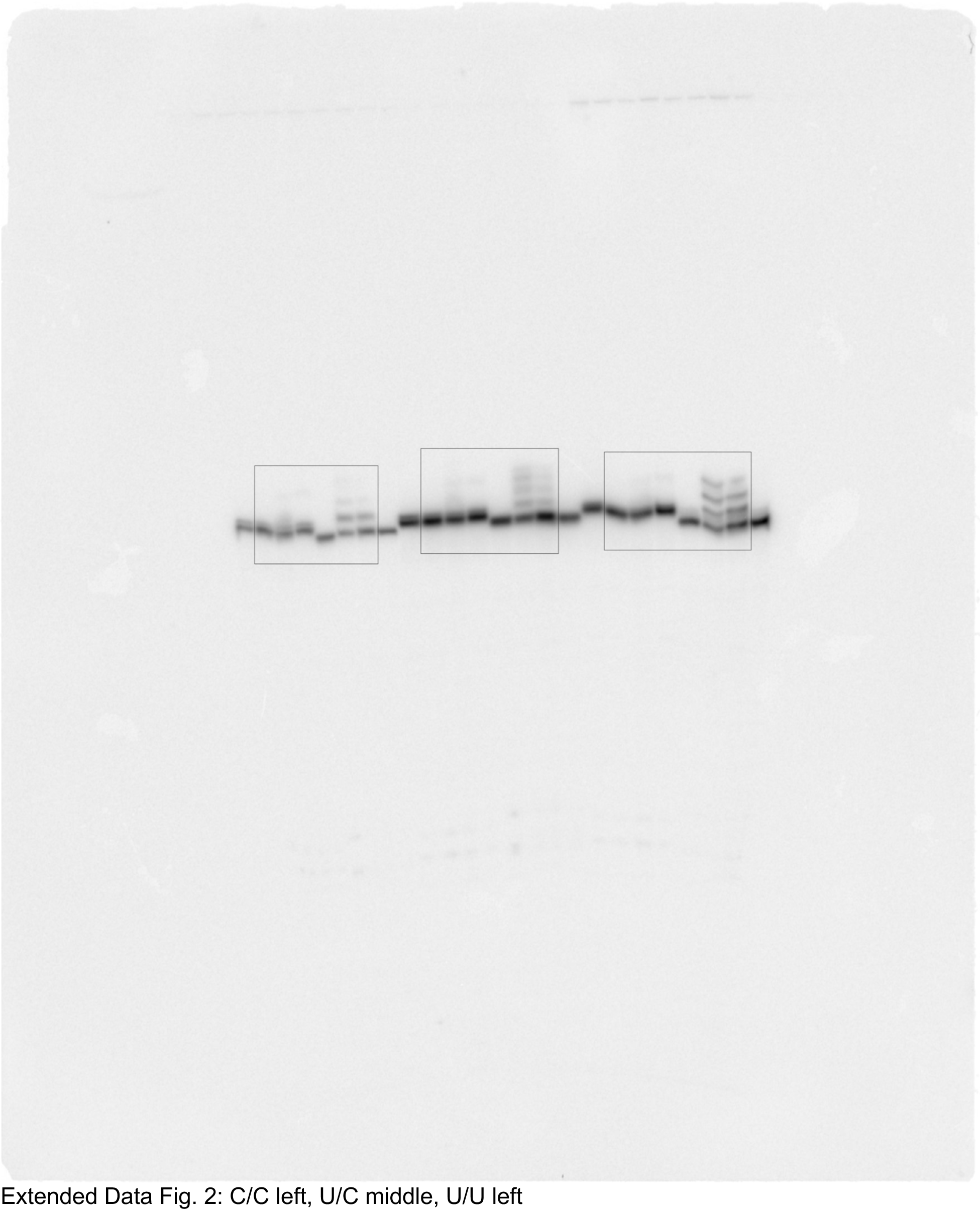

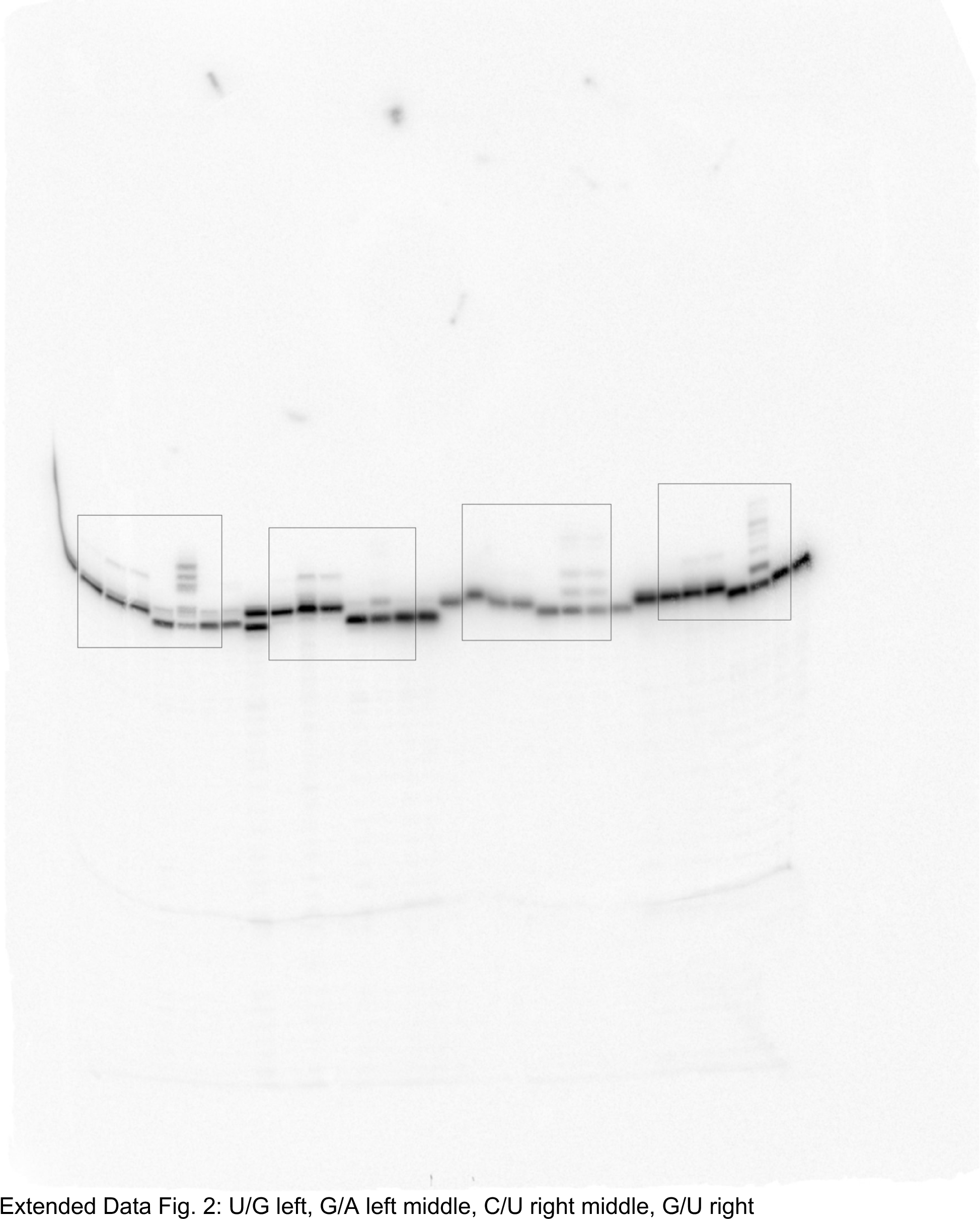

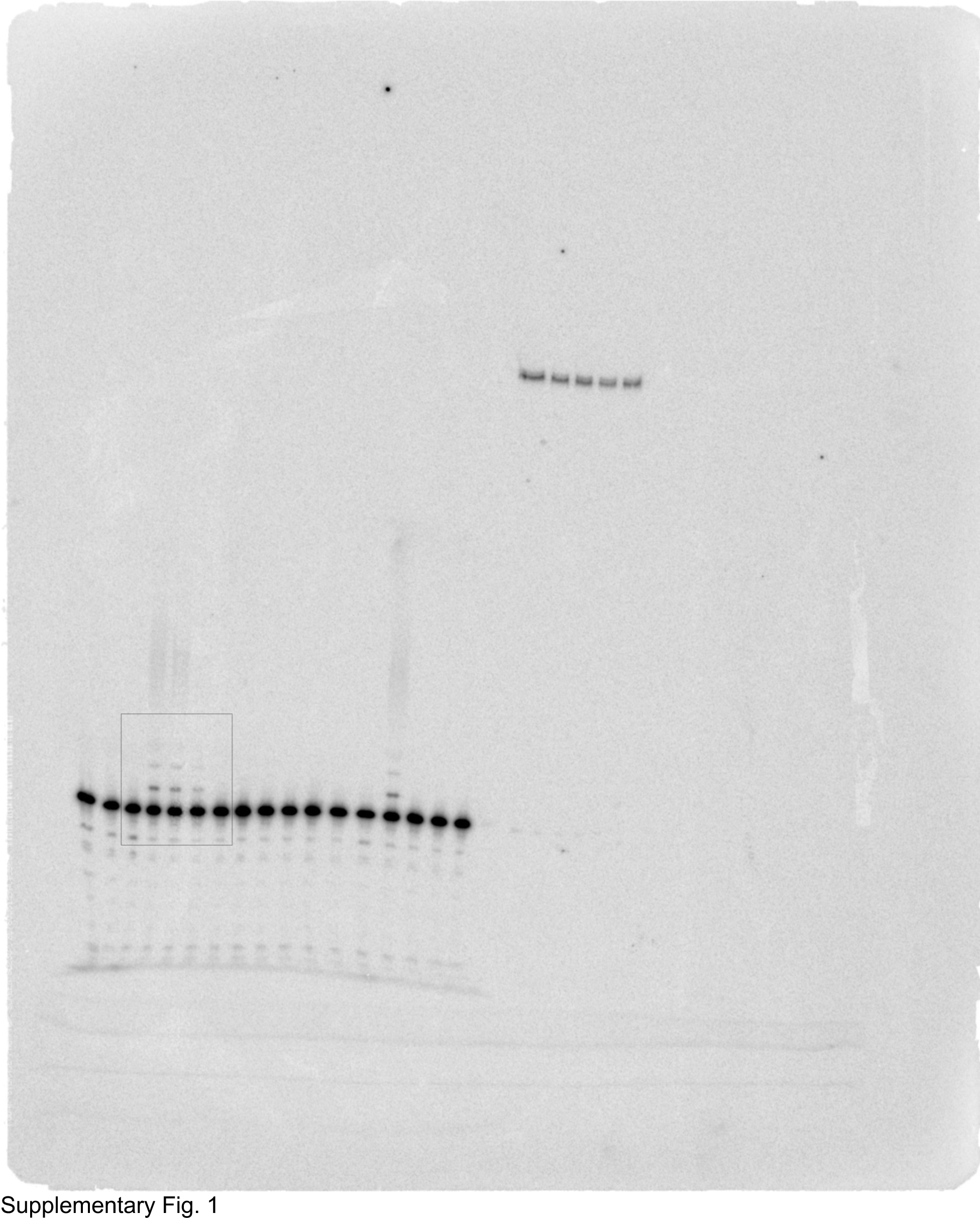

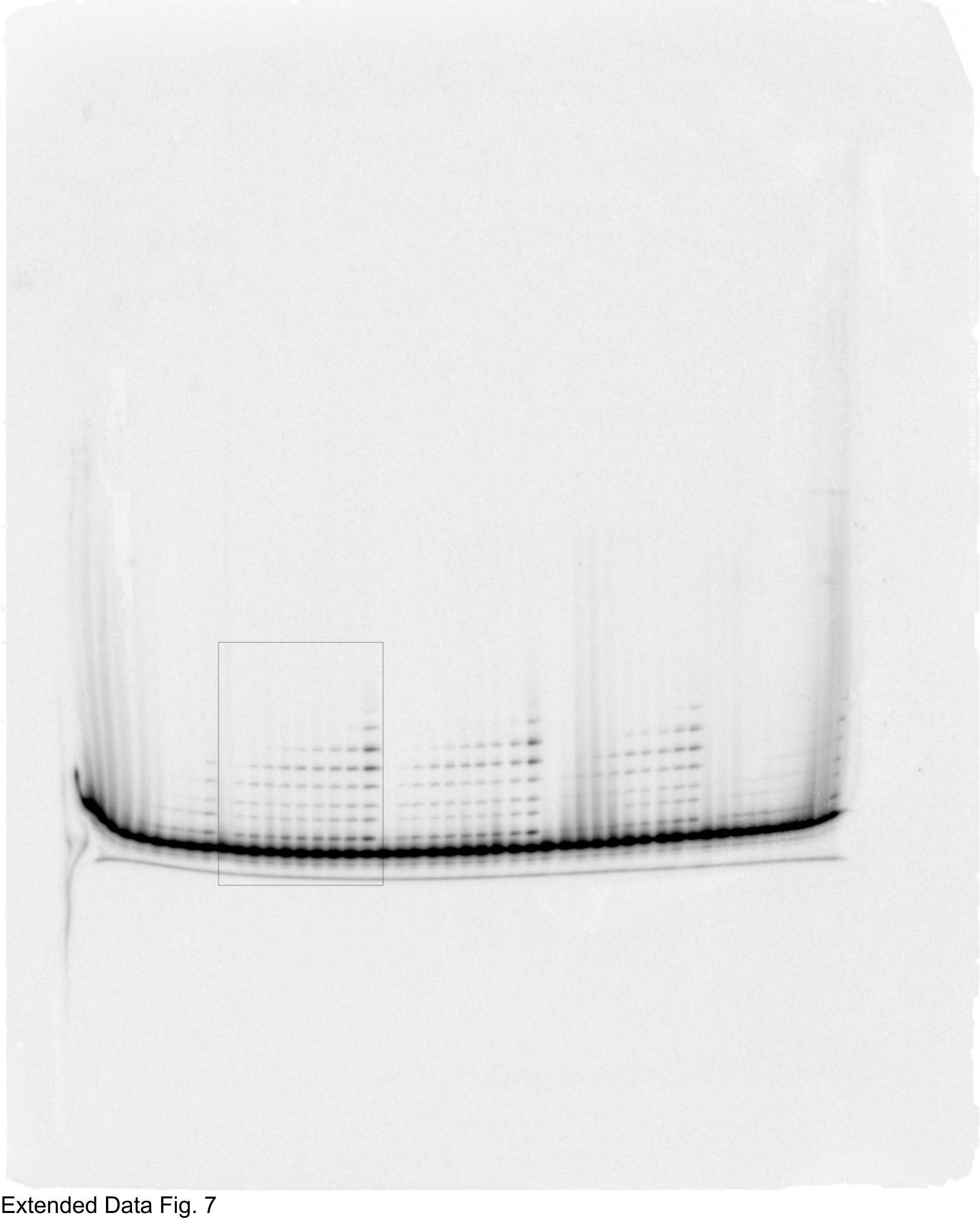

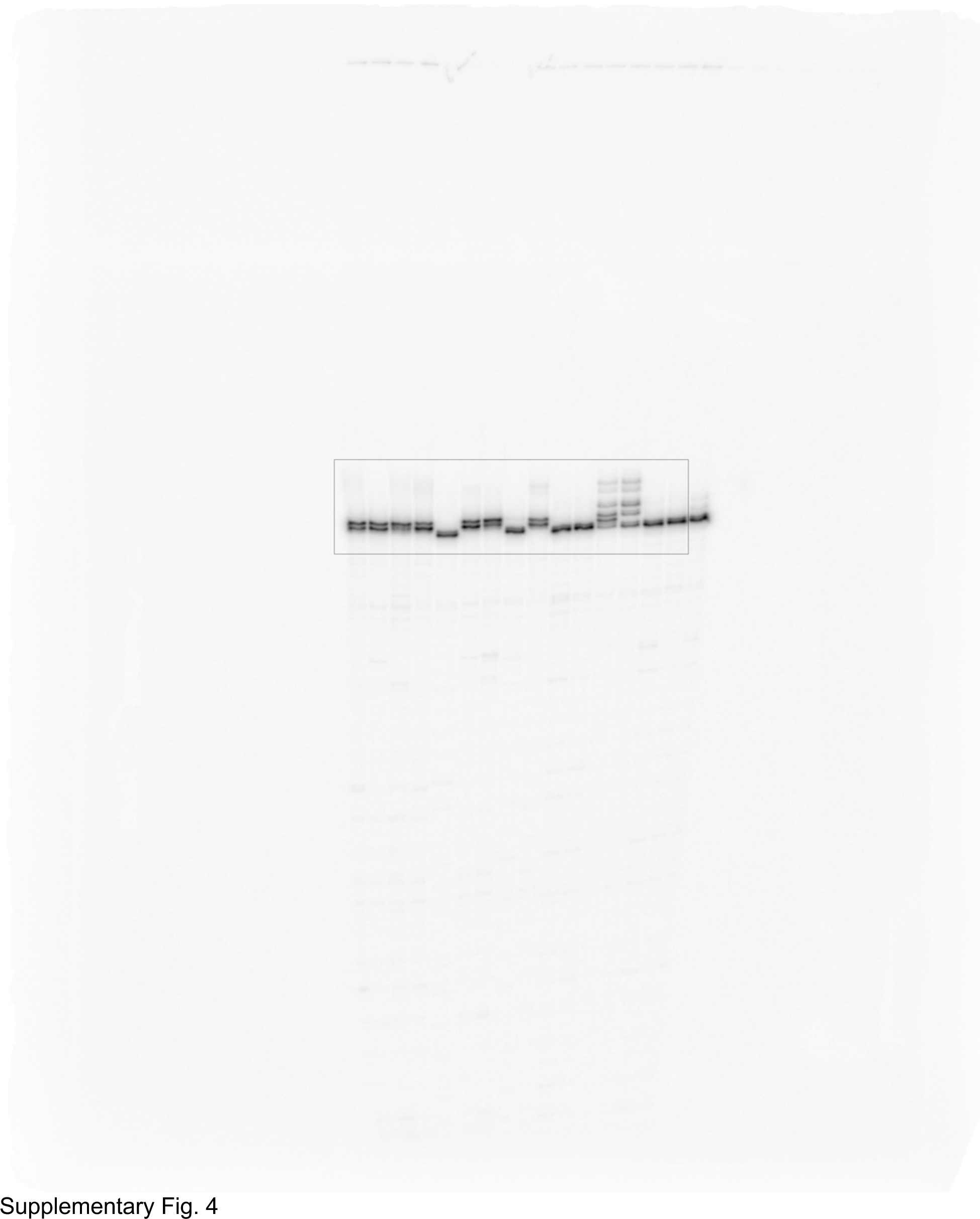

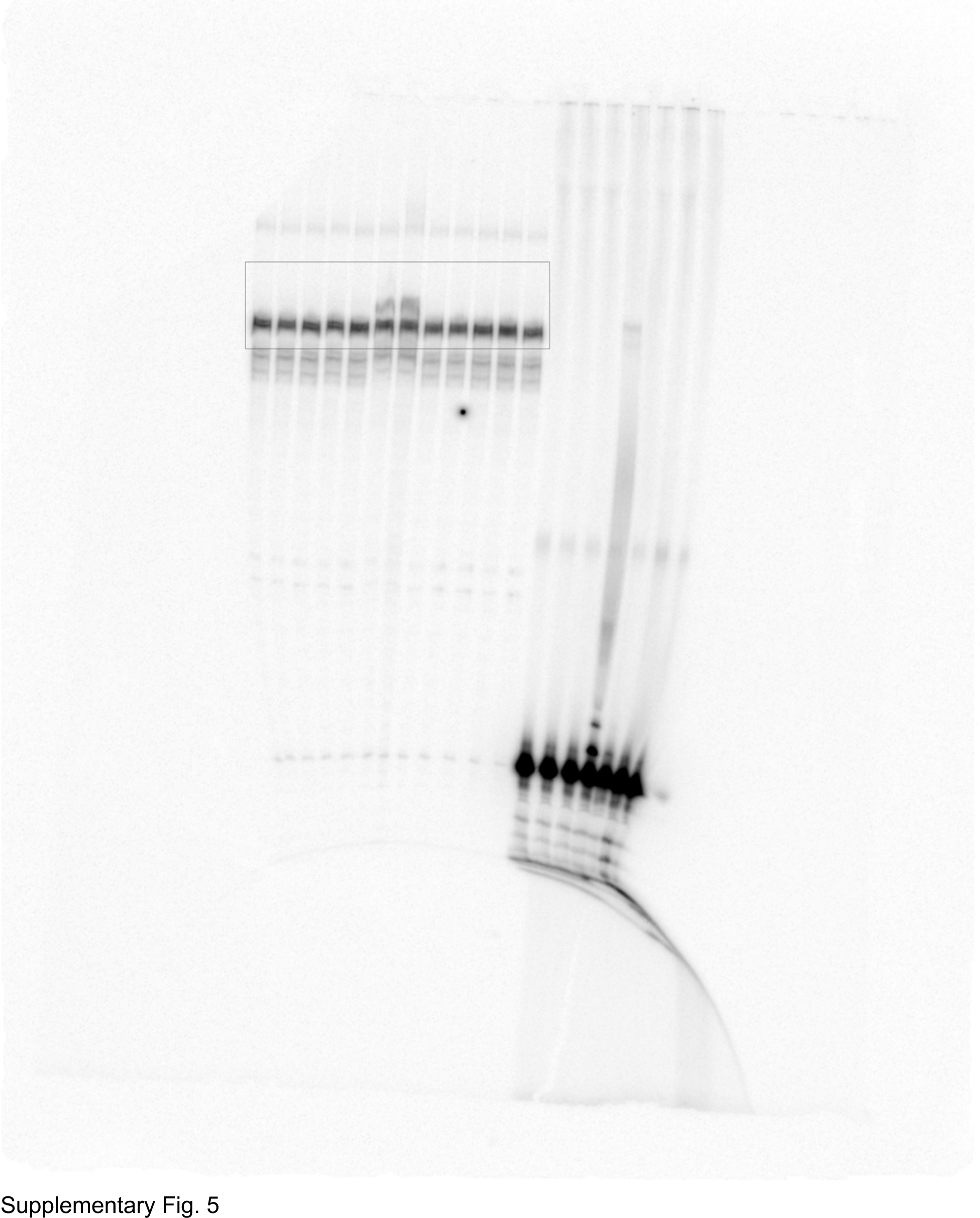

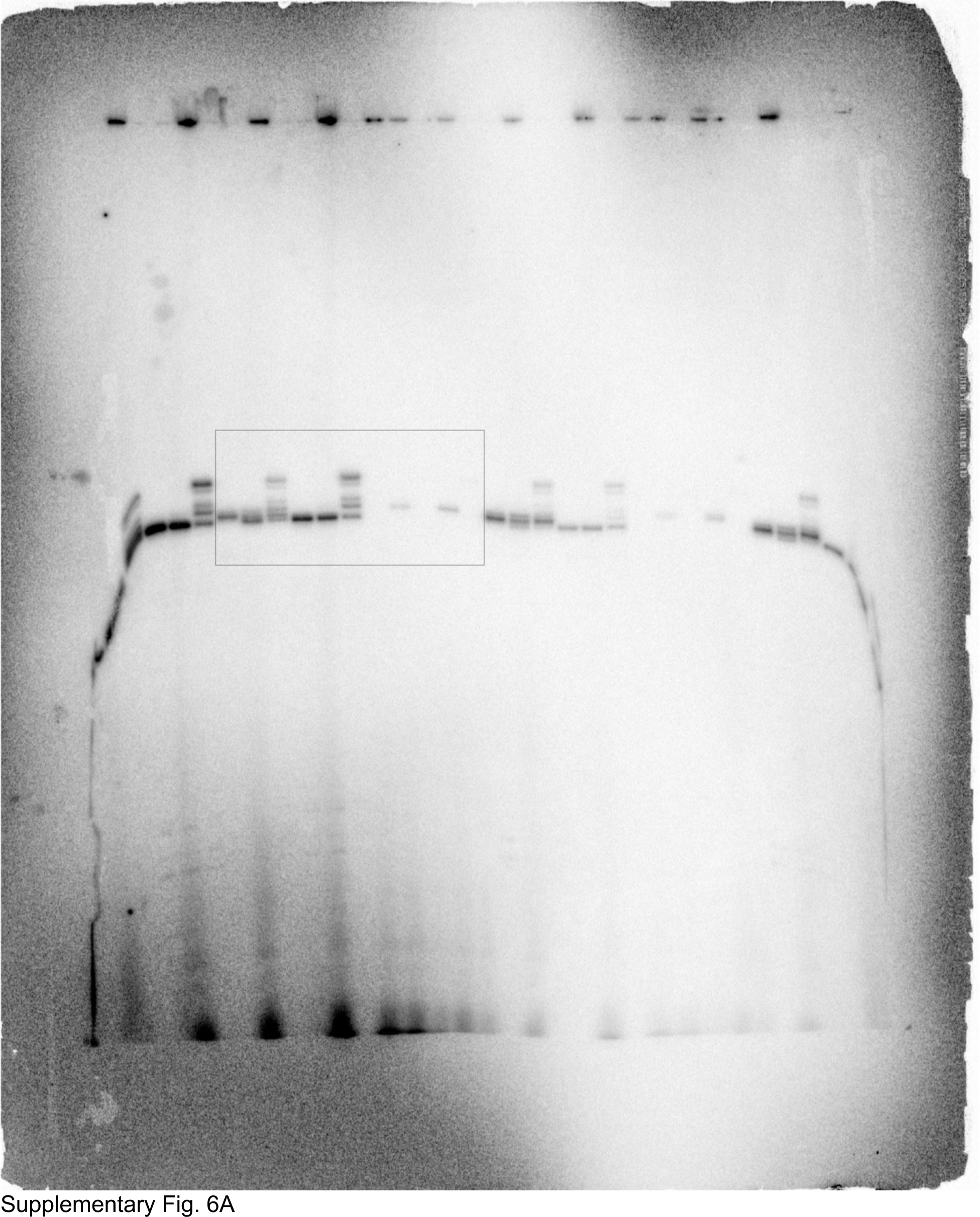

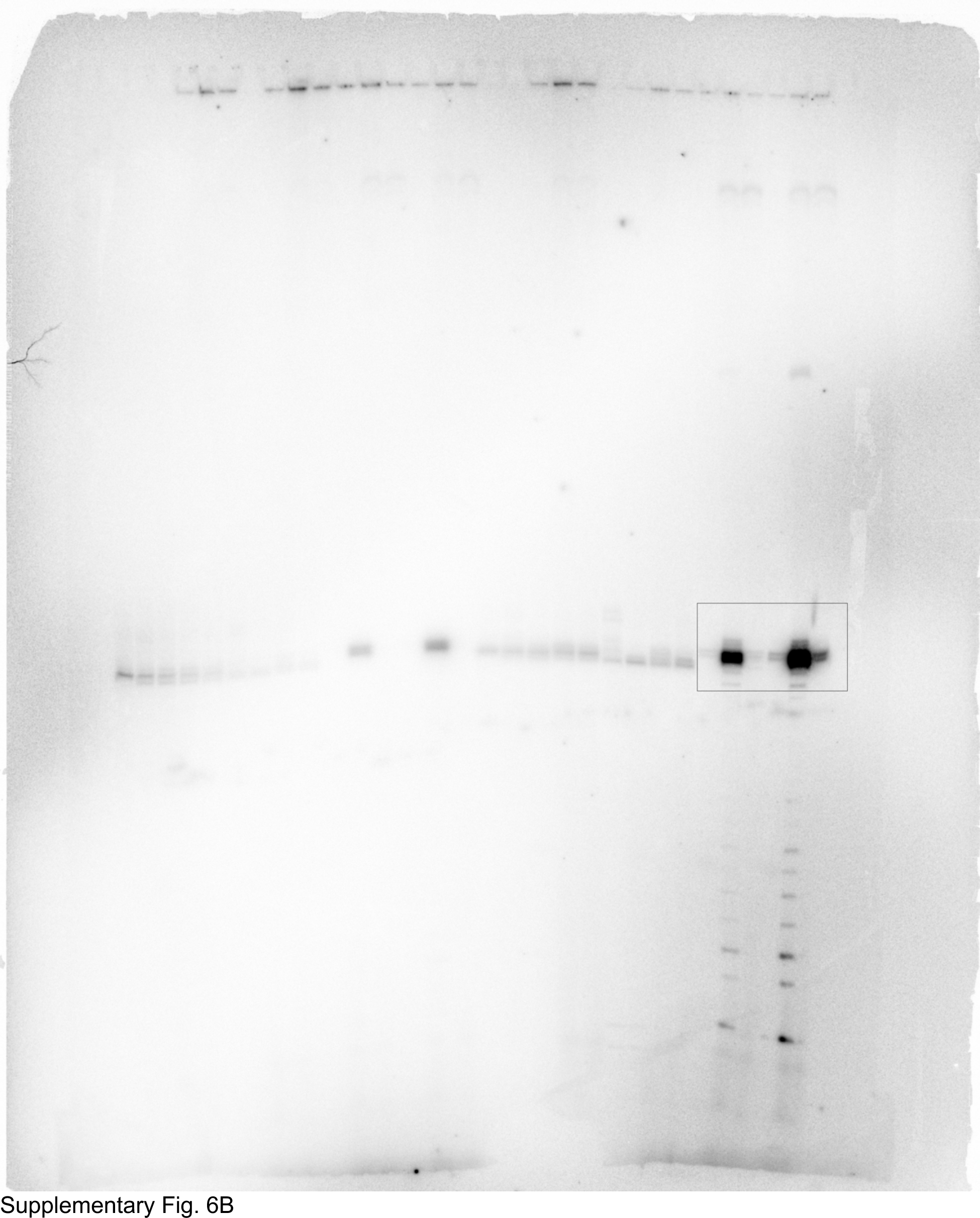

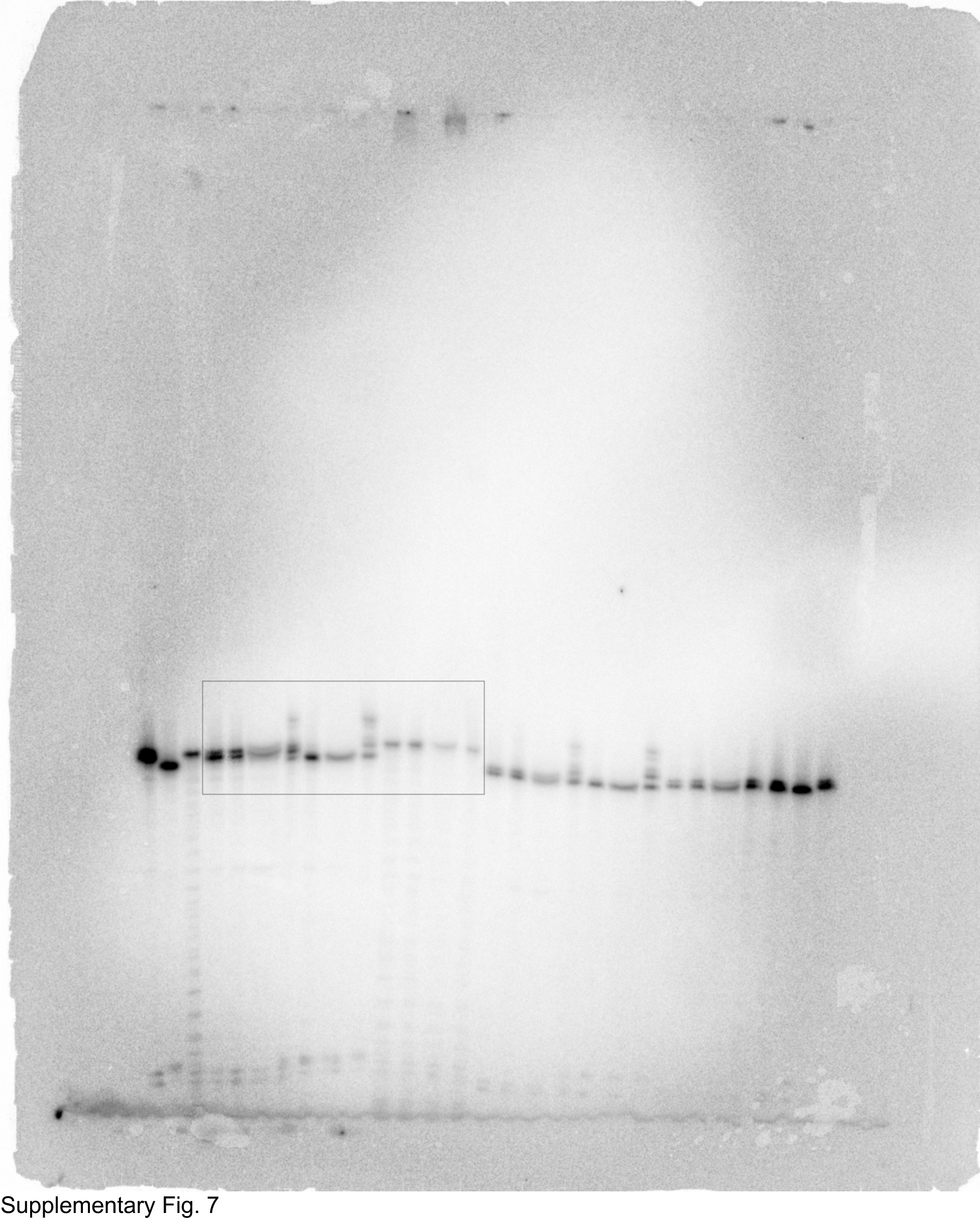

